# Multi-omics profiling identifies a metabolic and energetic signature associated with neuronal *dENL/AF9* suppression in aged flies

**DOI:** 10.64898/2026.09.30.755827

**Authors:** Ranchana Yeewa, Pitiporn Noisagul, Sivamoke Dissook, Chansunee Panto, Puttachat Poound, Salinee Jantrapirom, Luca Lo Piccolo

**Author notes:** Department of Biochemistry, Faculty of Medicine, Chiang Mai University, Chiang Mai, Thailand (Ranchana Yeewa).

## Abstract

Maintenance of neuronal function during aging requires metabolic adaptation, yet how metabolite-sensitive chromatin regulators contribute to this process remains unclear. YEATS-domain proteins, including ENL and AF9, recognize histone acylation marks and are positioned at the interface between cellular metabolism and transcriptional regulation. We previously showed that pan-neuronal suppression of *dENL/AF9*, the single *Drosophila* orthologue of mammalian ENL and AF9, extends lifespan, preserves healthspan, and enhances oxidative-stress resistance. Here, we used multi-omics and biochemical analyses to define the molecular state associated with neuronal *dENL/AF9* suppression. Transcriptomic profiling revealed a markedly stronger response in aged than young male fly heads, with prominent enrichment of fatty-acid degradation and several intermediary metabolic pathways in the aged dataset. Guided by this age-related transcriptional response, we focused subsequent profiling on aged heads, where independent molecular layers converged on a distinct metabolic and energetic signature. Lipidomics identified reduced abundance of multiple triglyceride-related and fatty acid derivatives, while metabolomics revealed changes in central-carbon/TCA-, nicotinamide-, and purine-associated features. Proteomics implicated mitochondrial and ATP-associated processes, accompanied by increased ATP abundance and a higher NAD/NADH ratio. Matched lipidome-metabolome integration further identified citrate as a cross-omics feature inversely associated with altered triglyceride and ceramide species, linking central-carbon and lipid-associated components of the molecular signature. Together, these findings identify a stronger transcriptional response to neuronal *dENL/AF9* suppression in aged flies and a distinct metabolic and energetic molecular signature in aged fly heads, providing candidate molecular correlates of the previously established longevity phenotype and a framework for studies of neuronal aging biology.

**Graphical abstract:** 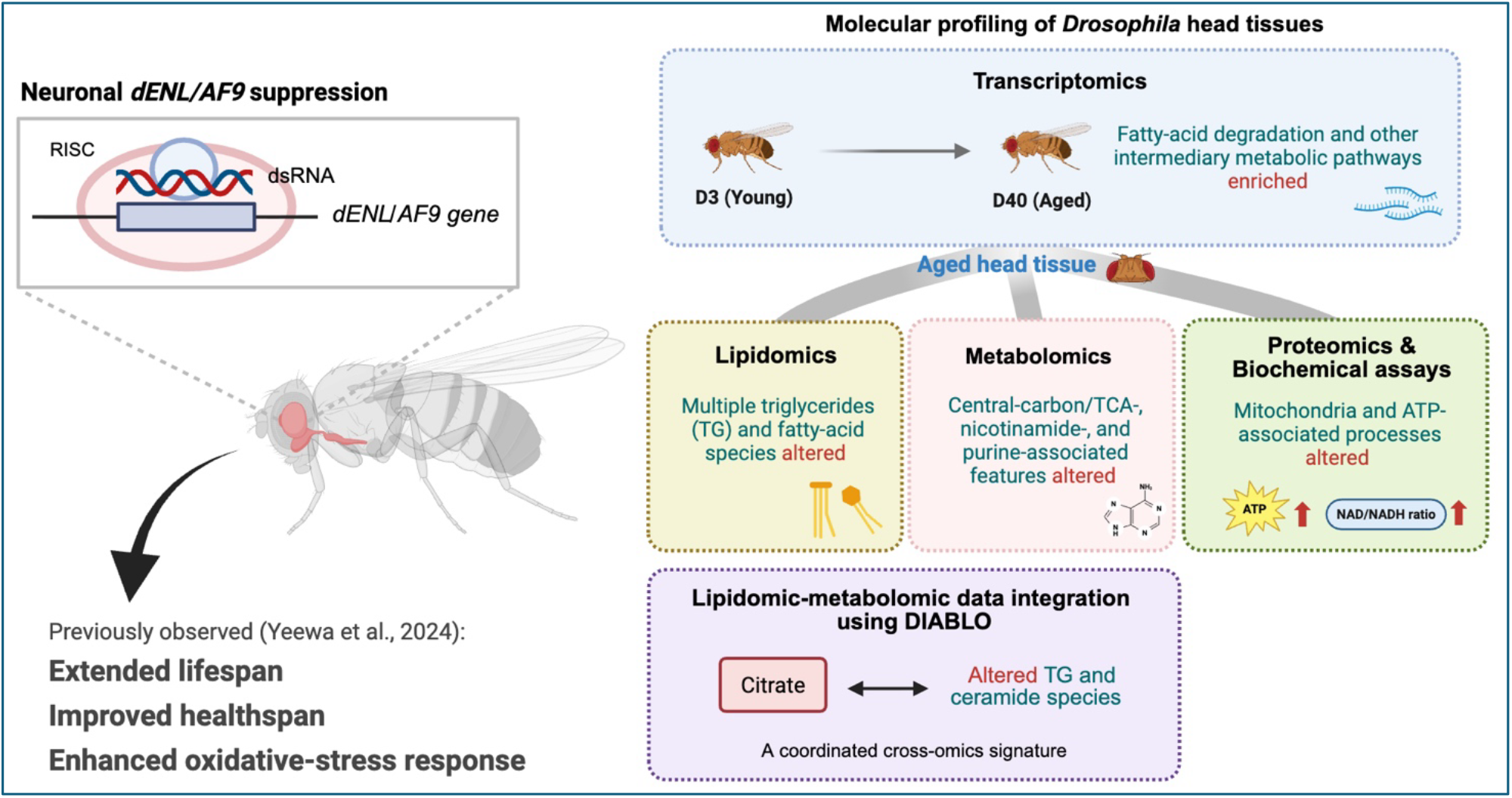

## Introduction

Maintenance of metabolic homeostasis becomes increasingly challenging in the aging nervous system. Neurons have sustained energetic demands and limited tolerance to disturbances in mitochondrial function, redox balance, and substrate availability, making metabolic resilience important for preserving neuronal function across the lifespan [1, 2]. Metabolism and chromatin regulation are closely interconnected: metabolites provide substrates and cofactors for chromatin-modifying reactions, while metabolite-sensitive chromatin mechanisms can translate changes in cellular metabolic state into transcriptional responses [3]. This coupling provides a molecular interface through which metabolic conditions can influence gene regulation and cellular adaptation. Understanding how such mechanisms operate in neurons is therefore particularly relevant to defining the molecular responses that support nervous-system function during aging.

YEATS (Yaf9, ENL, AF9, Taf14, Sas5) domain-containing proteins are evolutionarily conserved chromatin readers that recognize acylated lysine residues, including histone acetylation and crotonylation [4–6]. Because these histone modifications depend on metabolically derived acyl donors, YEATS-domain proteins are positioned at an interface between cellular metabolism and transcriptional regulation. Functional evidence for this relationship spans multiple biological systems. In yeast, Taf14 recognizes metabolically responsive histone crotonylation and contributes to transcriptional adaptation during metabolic cycling, whereas Yaf9 recognizes acetylated histones and coordinates recruitment of chromatin-regulatory factors during the oxidative phase of the metabolic cycle [7, 8]. In mammalian cells, AF9 regulates the gluconeogenic genes PCK2 and FBP1 through H3K9ac-dependent chromatin association and has also been linked to transcriptional regulation of purine metabolism through recognition of acetylated STAT6 [9, 10]. Our recent neuronal studies further extend the metabolic and homeostatic functions of YEATS-domain proteins. Pan-neuronal depletion of the *Drosophila* YEATS2 orthologue produces coordinated transcriptional changes, including downregulation of fatty-acid-oxidation and branched-chain-amino-acid-degradation programs [11]. Moreover, pharmacological inhibition of the ENL/AF9 YEATS domain alters PERK-dependent integrated stress response (ISR) signaling in neuronal models, with beneficial or detrimental effects depending on the prevailing stress context [12]. Because the ISR links stress-dependent translational control with transcriptional programs that regulate amino-acid metabolism, redox balance, and cellular homeostasis, these findings further connect YEATS-domain activity with neuronal homeostatic regulation. Together, current evidence indicates that YEATS readers can couple chromatin state to metabolic and stress-responsive programs, but how neuronal ENL/AF9 relates to the molecular state associated with physiological aging and longevity remains incompletely understood.

*Drosophila melanogaster* provides a genetically tractable system in which to address this question because the closely related mammalian ENL and AF9 proteins are represented by a single *Drosophila* orthologue, ear, hereafter referred to as *dENL/AF9* [13–15]. We previously demonstrated that pan-neuronal suppression of *dENL/AF9* extends lifespan and preserves multiple aspects of healthspan, including locomotor performance and sleep–wake organization [16]. Neuronal *dENL/AF9* suppression also increases resistance to oxidative stress, enhances catalase expression and activity, and reduces lipid peroxidation, without detectable changes in food intake or body weight [16]. These findings established neuronal *dENL/AF9* as a regulator of longevity and stress resilience; however, the molecular state associated with this phenotype remained unresolved. In particular, whether the transcriptional response to *dENL/AF9* suppression differs with age and whether the longevity-associated phenotype is accompanied by coordinated metabolic, energetic, and redox changes were unknown.

Here, we combined transcriptomic, lipidomic, metabolomic, proteomic, and biochemical analyses to characterize the molecular changes associated with neuronal *dENL/AF9* suppression. Transcriptomic profiling of young and aged fly heads was used to determine whether the transcriptional response to *dENL/AF9* suppression differed with age, whereas subsequent molecular profiling of aged fly heads was used to define the metabolic, energetic, and redox features associated with the previously established longevity phenotype. By integrating these complementary datasets, we further examined whether independently measured molecular layers converged on shared biological processes and whether matched lipidomic and metabolomic profiles formed a coordinated genotype-associated feature structure. Our findings identify an altered age-dependent transcriptional response together with a distinct metabolic, energetic, and redox molecular state associated with neuronal *dENL/AF9* suppression in aged fly heads.

## Methods

### Drosophila stocks, genetic crosses, and husbandry

*Drosophila melanogaster* were maintained on standard cornmeal–yeast–glucose medium at 25 °C and 55% relative humidity under a 12-h light/12-h dark cycle. Flies were maintained at low density, with approximately 15-20 adults per vial, and transferred to fresh food every 3 days.

Pan-neuronal RNA interference (RNAi) was achieved using the GAL4/UAS system. The neuronal synaptobrevin driver *nSyb-GAL4* (Bloomington Drosophila Stock Center [BDSC] #68222) was used to express double-stranded RNA in post-mitotic neurons. The RNAi lines used were UAS-*luciferase* RNAi (BDSC #31603), designated NTC IR and used as the principal non-targeting control throughout the omics experiments; UAS-*mCherry* RNAi (BDSC #35785), designated NTC IR#2 and used as an additional independent control in selected biochemical experiments; and UAS-*dENL/AF9* RNAi (BDSC #34798), designated dENL/AF9 IR. Prior to the experiments, the fly lines were backcrossed for six successive generations to the same *w* genetic background to minimize differences attributable to genetic background.

Experimental male progeny were generated by crossing the corresponding UAS-RNAi lines with *nSyb-GAL4*. Unless otherwise specified, comparisons throughout the study were performed between *nSyb>Luciferase RNAi* (NTC IR) and *nSyb>dENL/AF9 RNAi* (dENL/AF9 IR) flies.

### Experimental design and sample collection

The study was designed to characterize age-dependent molecular changes associated with pan-neuronal suppression of *dENL/AF9*. Adult male flies were collected at 3 days after eclosion (D3; young) and 40 days after eclosion (D40; aged). Transcriptomic profiling was performed at both D3 and D40 to determine whether the molecular response to *dENL/AF9* suppression differed with age. Subsequent lipidomic, metabolomic, proteomic, and biochemical analyses focused on D40 flies, where the transcriptomic response was most pronounced. The Ethics Committee of the Faculty of Medicine, Chiang Mai University, Thailand, approved the experimental protocol (Protocol code: 2569/INS-11).

For RNA-seq, three independent biological replicates were analyzed per genotype at each age, with each biological replicate consisting of 50 pooled fly heads. For untargeted lipidomic and metabolomic analyses, five independent biological replicates per genotype were analyzed at D40, with 50 pooled heads per replicate; lipidomic and metabolomic measurements were obtained from the same biological samples. Five biological replicates per genotype were initially prepared for proteomic profiling. Following quality assessment, one replicate per each group was excluded from the downstream proteomic statistical analysis because it was identified as an outlier, resulting in four biological replicates per genotype in the final proteomic dataset. The criterion and supporting quality-control evidence for this exclusion are described in the proteomics subsection.

For each experiment, flies were collected into 1.5-mL tubes, rapidly frozen in liquid nitrogen, and stored at −80 °C until sample preparation. Heads were separated from frozen flies before molecular extraction. Independent fly cohorts were used for transcriptomic and proteomic analyses, whereas the lipidomic and metabolomic datasets were derived from matched head samples. Additional independent cohorts were used for ATP and NAD/NADH measurements, as described below.

### RNA sequencing and transcriptomic analysis

#### RNA isolation, library preparation, and sequencing

For transcriptomic profiling, D3 and D40 male NTC IR and dENL/AF9 IR flies were collected and immediately frozen in liquid nitrogen. Three biological replicates were analyzed for each genotype at each age, with 50 pooled heads per replicate. Total RNA was isolated using the RNeasy Mini Kit (Qiagen, Valencia, CA, USA) according to the manufacturer’s instructions. RNA samples were maintained in GenTegra tubes and submitted to Macrogen, Inc. (Seoul, Republic of Korea) for library preparation and sequencing. RNA quality was evaluated using an Agilent 2100 Bioanalyzer. Samples with an RNA integrity number (RIN) >7 and an rRNA ratio >1 were used for library preparation. Strand-specific mRNA libraries were generated using the TruSeq Stranded mRNA Library Prep Kit and sequenced on an Illumina NovaSeq 6000 platform using 100-bp paired-end reads, yielding approximately 30 million reads per sample.

#### RNA-seq preprocessing and differential expression analysis

Raw sequencing files were retrieved from the sequencing provider and processed on the MedCMU high-performance computing platform – MedCMU HPC – using the nf-core/rnaseq pipeline v3.14.0 implemented through Nextflow [17, 18], following the computational workflow previously applied by our group [11]. Briefly, quality assessment of raw sequencing reads was performed using FastQC v0.12.0, and adapter and low-quality sequence trimming was performed using Trim Galore! v0.6.7 [19]. Putative contaminating genomic sequences were screened using BBSplit [20], and residual ribosomal RNA reads were filtered in silico using SortMeRNA [21]. Processed reads were aligned to the BDGP6.46 assembly (Ensembl release 112) using HISAT2 v2.2.1 with reverse-stranded settings [22, 23]. Alignment files were coordinate-sorted and indexed using SAMtools v1.17 [24]. Duplicate molecules in UMI-tagged libraries were collapsed with UMI-tools [25], while duplicates in non-UMI libraries were marked with Picard MarkDuplicates v3.0.0. Quality-control metrics generated throughout the pipeline were compiled using MultiQC v1.19, as summarized in Supplementary Table 1. Reference-guided transcript quantification was performed using StringTie2 v2.2.1 with Ensembl release 112 annotation [26]. Gene-level count matrices were subsequently analyzed using DESeq2 in R v4.1 [27]. Differentially expressed genes (DEGs) between dENL/AF9 IR and age-matched NTC IR samples were defined using an absolute fold-change threshold of ≥1.5 and a Benjamini–Hochberg-adjusted p-value <0.05. A total of 16,885 genes were detected across the transcriptomic dataset.

#### Gene set enrichment and leading-edge analyses

Pathway-level transcriptional changes were evaluated using Gene Set Enrichment Analysis (GSEA) v4.3.2, using previously described parameters [11]. DESeq2-normalized gene-expression values were analyzed against a predefined collection of 137 Kyoto Encyclopedia of Genes and Genomes (KEGG) pathways for *D. melanogaster* [28]. Separate enrichment analyses were performed for D3 and D40 dENL/AF9 IR samples relative to their respective age-matched NTC IR controls. GSEA was performed using gene-set permutation, the Signal2Noise ranking metric, and weighted enrichment scoring based on the Kolmogorov–Smirnov statistic. GSEA pathways with a false discovery rate (FDR) q<0.25 were considered enriched according to the conventional GSEA discovery threshold [29], whereas FDR q<0.05 was considered stronger evidence of enrichment. Pathways with q values between 0.05 and 0.25 were interpreted cautiously as exploratory. Normalized enrichment scores (NESs), nominal P values, and FDR q-values were used to summarize pathway-level changes. Leading-edge analysis was subsequently performed for selected enriched pathways to identify the subset of genes contributing most strongly to the enrichment signal. Heatmaps of leading-edge genes and summary bubble plots of enriched pathways were generated using the SRplot online visualization platform [30].

### Untargeted lipidomics analysis

#### Sample preparation and lipid extraction

Untargeted lipidomic analysis was performed by Baiya Phytopharm Co., Ltd., Chulalongkorn University, Thailand. Five independent biological replicates of D40 NTC IR and dENL/AF9 IR fly heads were analyzed, with each replicate containing 50 pooled heads. Lipidomic and metabolomic measurements were performed using the same biological samples. Samples were extracted with 70% methanol containing 50 ng/mL sulfadimethoxine as an internal standard to obtain a final sample concentration of 10 mg/mL. Following centrifugation at 10,000 rpm for 10 min, the supernatant was collected for metabolomic and lipidomic analyses. An aliquot of 300 µL supernatant was mixed with 210 µL chloroform and 5 µL of a 100 µg/mL internal-standard solution containing tolnaftate and natamycin, followed by centrifugation at 10,000 rpm for 10 min. The lower organic phase was collected for lipidomic analysis, dried, and reconstituted in 200 µL isopropanol/acetonitrile/water (2:1:1, v/v/v). A pooled quality-control (QC) sample was prepared by combining 50 µL from each biological sample. Serial dilutions of the pooled QC sample corresponding to 100%, 80%, 50%, 20%, 10%, 1%, and 0% were prepared to evaluate analytical response and feature reproducibility. Prior to LC–MS analysis, samples were centrifuged at 14,000 rpm for 10 min, and the resulting supernatants were transferred to LC–MS vials.

#### LC–QTOF-MS acquisition

Lipidomic profiling was performed using an Agilent 6545XT LC/Q-TOF mass spectrometer operated in high-resolution positive- and negative-ionization modes. Chromatographic separation was achieved using a Poroshell 120 EC-C18 column (2.1 × 100 mm, 2.7 µm) maintained at 55 °C, with an injection volume of 10 µL and a flow rate of 0.3 mL/min. Mobile phase A consisted of water/acetonitrile/isopropanol (2:1:1, v/v/v) containing 10 mM ammonium formate, whereas mobile phase B consisted of isopropanol/acetonitrile (9:1, v/v) containing 10 mM ammonium formate. The gradient was programmed as follows: 0–0.5 min, 100% A; 2.0 min, 60% A/40% B; 14.0 min, 5% A/95% B; 14.5–16.5 min, 100% B; and 17.0–19.0 min, 100% A for column re-equilibration. The mass spectrometer was operated with a drying-gas temperature of 325 °C, drying-gas flow of 13 L/h, sheath-gas temperature of 275 °C, sheath-gas flow of 12 L/h, and nebulizer pressure of 45 psi. Capillary voltages were 4,000 V in positive mode and 3,000 V in negative mode. MS1 spectra were acquired over *m/z* 40–1,700 and MS/MS spectra over *m/z* 25–1,000. Collision energies were 20 eV in positive mode and 10 eV in negative mode. Reference ions at *m/z* 121.0509 and 922.0098 in positive mode and *m/z* 112.9856 and 1033.9881 in negative mode were used for mass correction.

#### Lipidomic data processing and statistical analysis

Raw LC–MS data were processed using MS-DIAL v5.3. Features were aligned against a pooled QC sample and normalized using locally weighted scatterplot smoothing (LOWESS) together with the internal standards tolnaftate for positive-ion data and natamycin for negative-ion data. Lipid annotations were assigned by comparison with the MS-DIAL ESI positive/negative MS/MS libraries derived from authentic standards and the MS-DIAL Tandem Mass Spectral Atlas. Features were retained if they eluted between 0.2 and 18 min, showed a Pearson correlation coefficient ≥0.70 across the QC dilution series, exhibited a coefficient of variation <30% among pooled QC samples, and had an identification score ≥0.70.

The processed feature-intensity matrix was subsequently analyzed using MetaboAnalyst 6.0. Because the input dataset had already undergone analytical normalization, no additional sample normalization was applied. Variables were Pareto-scaled before multivariate analysis. Principal component analysis (PCA) was used to assess overall variation among samples, whereas partial least-squares discriminant analysis (PLS-DA) was used to evaluate genotype-associated separation. PLS-DA model performance was evaluated using five-fold cross-validation in MetaboAnalyst 6.0. Up to five latent components were assessed, and the optimal number of components was selected based on the cross-validated Q² value. R² and Q² were used to describe model fit and predictive performance, respectively. Variable importance in projection (VIP) scores were obtained from the PLS-DA model, and lipid features with VIP scores >1 were considered major contributors to group discrimination and were retained for subsequent visualization and biological interpretation. Hierarchical clustering and heatmap visualization were performed on selected lipid features.

### Untargeted metabolomics analysis

#### Metabolite extraction and LC–QTOF-MS acquisition

Untargeted metabolomic profiling was performed on the same D40 head extracts used for lipidomic analysis. Following methanol extraction and chloroform partitioning as described above, the upper aqueous phase was collected for metabolomic analysis. A pooled QC sample and serial QC dilution series were prepared as described for the lipidomic workflow. Samples were centrifuged at 14,000 rpm for 10 min prior to transfer into LC–MS vials. Chromatographic separation was performed using a Poroshell 120 EC-C18 column (2.1 × 100 mm, 2.7 µm) maintained at 50 °C, with an injection volume of 10 µL and a flow rate of 0.4 mL/min. Mobile phase A consisted of 0.1% formic acid in water and mobile phase B consisted of 0.1% formic acid in acetonitrile. The gradient was programmed as follows: 0–0.5 min, 100% A; 10.5 min, 45% A/55% B; 12.5 min, 25% A/75% B; 14.0–17.0 min, 100% B; and 17.5–20.0 min, 100% A for re-equilibration. Mass spectrometric analysis was performed on an Agilent 6545XT LC/Q-TOF system in high-resolution positive- and negative-ionization modes using the same source conditions described for lipidomics. MS1 and MS/MS acquisition ranges were *m/z* 40–1,700 and *m/z* 25–1,000, respectively. Collision energies were 20 eV in positive mode and 10 eV in negative mode. Reference ions at *m/z* 121.0509 and 922.0098 in positive mode and *m/z* 112.9856 and 1033.9881 in negative mode were used for mass correction.

#### Metabolomic data processing and feature annotation

Raw metabolomic data were processed using MS-DIAL v5.3 and aligned against a pooled QC sample. Feature intensities were normalized using LOWESS and the sulfadimethoxine internal standard. Features were retained if they met the following criteria: retention time 0.2–18 min, Pearson correlation coefficient ≥0.70 across the QC dilution series, QC coefficient of variation <30%, provider-specific identification score ≥0.70, and mass error within ±20 ppm.

Metabolite annotations were assigned by the analytical provider using MS-DIAL database matching and were accompanied by a provider-specific Total score, representing relative annotation confidence, with higher scores indicating greater confidence. The Total score was not interpreted as a probability or converted to a formal Metabolomics Standards Initiative (MSI) identification level. In the provider output, annotations were further designated as either “low score”, indicating limited-confidence database annotation, or “no MS2”, indicating annotation without supporting MS/MS information. Because none of the metabolite identities were independently confirmed against authentic reference standards in this study, all named metabolites were treated as putatively annotated features, and biological interpretation emphasized coherent pathway-level patterns rather than definitive identification of individual compounds.

#### Multivariate and pathway analyses

The processed metabolite-intensity matrix was analyzed using MetaboAnalyst 6.0. No additional sample normalization was applied to the analytically normalized input data, and variables were Pareto-scaled before statistical analysis. PCA and PLS-DA were performed to assess overall metabolic variation and genotype-associated discrimination, respectively. PLS-DA model performance was evaluated using five-fold cross-validation in MetaboAnalyst 6.0. PLS-DA performance was assessed by cross-validation, with R² and Q² used to evaluate model fit and predictive ability, respectively. VIP scores were extracted from the PLS-DA model, and features with VIP scores >1 were retained as discriminatory metabolites for subsequent visualization and pathway analysis.

Putatively annotated VIP metabolites were subjected to pathway analysis using the *D. melanogaster* Kyoto Encyclopedia of Genes and Genomes (KEGG) metabolic pathway library implemented in MetaboAnalyst 6.0. Pathway enrichment was evaluated using the hypergeometric test, and pathway topology was assessed using relative-betweenness centrality. All compounds included in the selected pathway library were used as the reference metabolome. Pathways were visualized in a scatter plot according to enrichment significance and pathway impact. For graphical presentation, pathways with −log10(*P*) >0.5 or pathway impact >0.1 were highlighted. Raw *P* values, Holm-adjusted *P* values, false discovery rates, and pathway-impact values are reported in Table 1.

**Table 1.** Metabolic pathways associated with aged *dENL/AF9*-knockdown fly heads.

| KEGG pathway | Total | Expected | Hits | Raw p | Holm adjust | FDR | Impact |
| --- | --- | --- | --- | --- | --- | --- | --- |
| Nicotinate and nicotinamide metabolism | 9 | 0.229 | 3 | 0.001 | 0.086 | 0.086 | 0.113 |
| Purine metabolism | 68 | 1.729 | 6 | 0.006 | 0.426 | 0.216 | 0.136 |
| Citrate cycle (TCA cycle) | 20 | 0.508 | 3 | 0.013 | 0.940 | 0.322 | 0.169 |
| Glycerophospholipid metabolism | 32 | 0.814 | 3 | 0.044 | 1 | 0.856 | 0.154 |
| Phenylalanine, tyrosine and tryptophan biosynthesis | 4 | 0.102 | 1 | 0.098 | 1 | 1 | 0.500 |
| Alanine, aspartate and glutamate metabolism | 23 | 0.585 | 2 | 0.114 | 1 | 1 | 0.000 |
| One carbon pool by folate | 23 | 0.585 | 2 | 0.114 | 1 | 1 | 0.000 |
| Glycolysis or Gluconeogenesis | 26 | 0.661 | 2 | 0.139 | 1 | 1 | 0.179 |
| Phenylalanine metabolism | 7 | 0.178 | 1 | 0.165 | 1 | 1 | 0.379 |
| Glycine, serine and threonine metabolism | 29 | 0.737 | 2 | 0.166 | 1 | 1 | 0.000 |
| Arginine and proline metabolism | 29 | 0.737 | 2 | 0.166 | 1 | 1 | 0.178 |
| Valine, leucine and isoleucine biosynthesis | 8 | 0.203 | 1 | 0.187 | 1 | 1 | 0.000 |
| Arginine biosynthesis | 13 | 0.331 | 1 | 0.286 | 1 | 1 | 0.341 |
| Butanoate metabolism | 14 | 0.356 | 1 | 0.304 | 1 | 1 | 0.000 |
| Fructose and mannose metabolism | 16 | 0.407 | 1 | 0.340 | 1 | 1 | 0.023 |
| Propanoate metabolism | 20 | 0.508 | 1 | 0.405 | 1 | 1 | 0.000 |
| Pantothenate and CoA biosynthesis | 20 | 0.508 | 1 | 0.405 | 1 | 1 | 0.000 |
| Pentose phosphate pathway | 22 | 0.559 | 1 | 0.436 | 1 | 1 | 0.219 |
| Pyruvate metabolism | 23 | 0.585 | 1 | 0.450 | 1 | 1 | 0.147 |
| Glyoxylate and dicarboxylate metabolism | 24 | 0.610 | 1 | 0.465 | 1 | 1 | 0.110 |
| Glutathione metabolism | 26 | 0.661 | 1 | 0.492 | 1 | 1 | 0.018 |
| Lipoic acid metabolism | 27 | 0.686 | 1 | 0.505 | 1 | 1 | 0.000 |
| Sphingolipid metabolism | 27 | 0.686 | 1 | 0.505 | 1 | 1 | 0.004 |
| Inositol phosphate metabolism | 28 | 0.712 | 1 | 0.518 | 1 | 1 | 0.000 |
| Folate biosynthesis | 28 | 0.712 | 1 | 0.518 | 1 | 1 | 0.000 |
| Cysteine and methionine metabolism | 32 | 0.814 | 1 | 0.567 | 1 | 1 | 0.000 |
| Tyrosine metabolism | 33 | 0.839 | 1 | 0.578 | 1 | 1 | 0.000 |
| Valine, leucine and isoleucine degradation | 38 | 0.966 | 1 | 0.631 | 1 | 1 | 0.000 |
| Amino sugar and nucleotide sugar metabolism | 39 | 0.992 | 1 | 0.640 | 1 | 1 | 0.000 |
| Pyrimidine metabolism | 40 | 1.017 | 1 | 0.650 | 1 | 1 | 0.044 |

### Proteomic analysis

#### Protein extraction and enzymatic digestion

Untargeted proteomic profiling was performed by Baiya Phytopharm Co., Ltd., Chulalongkorn University, Thailand, using independent D40 NTC IR and dENL/AF9 IR head samples. Five biological replicates per genotype, each comprising 50 pooled heads, were initially subjected to protein extraction and LC–MS analysis. Head samples were homogenized in 6 M urea prepared in 50 mM Tris-HCl at a ratio of approximately 3 mg tissue per 100 µL lysis buffer. Two metal beads were added to each sample, followed by homogenization at 50 Hz for 3 min, repeated twice. Samples were centrifuged at 14,000 rpm for 10 min, and the supernatants were collected. Proteins were precipitated by addition of four volumes of ice-cold 100% methanol and recovered by centrifugation at 14,000 rpm for 10 min. Protein pellets were resuspended in 50 µL of 50 mM ammonium bicarbonate buffer, and protein concentrations were normalized to the lowest concentration among samples. For enzymatic digestion, 40 µL of normalized protein solution was reduced with 10 mM dithiothreitol at 65 °C for 30 min and subsequently alkylated with 25 mM iodoacetamide for 20 min at room temperature in the dark. Proteins were digested overnight at 37 °C following addition of 1.5 µL of 0.1 mg/mL trypsin. Digestion was terminated by addition of 10% formic acid. Samples were centrifuged at 14,000 rpm for 10 min, and the peptide-containing supernatants were transferred to LC– MS vials.

#### LC–QTOF-MS proteomic acquisition

Peptides were analyzed using an Agilent 6545XT LC/Q-TOF mass spectrometer coupled to an Agilent Peptide Map column (2.1 × 150 mm, 2.7 µm) maintained at 60 °C. An injection volume of 20 µL was used, with a total chromatographic run time of 85 min and a flow rate of 0.4 mL/min. Mobile phase A consisted of water containing 0.1% formic acid, and mobile phase B consisted of acetonitrile containing 0.1% formic acid. The gradient was programmed as follows: 0–2 min, 100% A; 35 min, 80% A/20% B; 55 min, 70% A/30% B; 65 min, 50% A/50% B; 70–75 min, 10% A/90% B; and 80–85 min, 100% A. Mass spectrometric acquisition was performed in positive-ion mode with a gas temperature of 325 °C, drying-gas flow of 13 L/h, nebulizer pressure of 35 psi, capillary voltage of 4,000 V, and nozzle voltage of 500 V. MS spectra were acquired over *m/z* 100– 1,700 and MS/MS spectra over *m/z* 25–1,000. The reference mass was *m/z* 922.0098.

#### Protein identification and label-free quantification

Agilent raw data files were converted to mzXML format using MSConvert and OpenMS and subsequently searched using MaxQuant v2.6.3 against the UniProt Drosophila melanogaster proteome database (taxonomy ID 7227). Trypsin was specified as the digestion enzyme. Carbamidomethylation of cysteine was defined as a fixed modification, whereas N-terminal acetylation and methionine oxidation were included as variable modifications. The minimum label-free quantification (LFQ) ratio count was set to 1. A discovery-oriented false discovery rate (FDR) threshold of 0.10 was applied at the protein and peptide-spectrum-match levels to increase proteome coverage. Protein groups identified only by site, reverse-sequence matches, and potential contaminants were removed. A total of 287 protein groups were retained for downstream analysis; of these, 229 protein groups had a MaxQuant protein-level q-value <0.01.

#### Proteomic statistical and functional analyses

LFQ intensities were subsequently analyzed using MetaboAnalyst 6.0. Features containing only zero values or a single non-zero value across the dataset were removed before statistical analysis. The processed data were Pareto-scaled prior to multivariate analysis. PCA was used to assess global sample variation, whereas PLS-DA was used to evaluate genotype-associated discrimination. PLS-DA performance was assessed by cross-validation, with R² and Q² used to evaluate model fit and predictive ability, respectively. Five biological replicates per genotype were initially analyzed. Unsupervised PCA and hierarchical clustering consistently identified one sample from each genotype as a marked outlier relative to the remaining biological replicates. These two samples were excluded before differential protein and functional enrichment analyses, resulting in a final sample size of four biological replicates per genotype. Differentially abundant proteins (DAPs) between dENL/AF9 IR and NTC IR samples were defined using an absolute fold-change threshold of ≥1.5 and an unadjusted *P* value <0.05. Functional enrichment of DAPs was evaluated using the Database for Annotation, Visualization and Integrated Discovery (DAVID), including Gene Ontology biological process, molecular function and cellular component categories, together with KEGG and Reactome pathway annotations. Protein–protein interaction relationships among DAPs were further examined using the STRING database to identify functionally connected protein clusters.

### ATP measurement

ATP levels in aged fly heads were measured using the ATP Assay Kit (MAK473; Sigma-Aldrich/Merck) according to the manufacturer’s protocol with modifications for *Drosophila* head tissue. Three experimental groups were assessed: the principal Luciferase RNAi control (NTC IR), an independent mCherry RNAi control (NTC IR#2), and dENL/AF9 IR. For each biological replicate, 30 pooled heads from D40 male flies were homogenized in cold phosphate-buffered saline (PBS). Tissue debris was removed by centrifugation, and the resulting supernatant was used for ATP determination. In parallel, an aliquot of the same extract was used to determine total protein concentration by the Bradford assay for normalization. An ATP standard curve ranging from 0 to 30 µM was prepared from the supplied ATP standard. Samples and standards were transferred to a white opaque 96-well plate, and ATP was quantified using a luciferase-based bioluminescence reaction in which ATP-dependent oxidation of D-luciferin generates a luminescent signal proportional to ATP concentration. The reaction mixture was added to each well, and relative luminescence units (RLU) were measured immediately using a microplate luminometer. ATP concentrations were calculated from blank-corrected luminescence values using the corresponding standard curve and normalized to protein concentration. The ATP calibration curve used for quantification is shown in Supplementary Fig. 8.

### NAD and NADH measurement

NAD and NADH levels were measured in aged fly heads using the NAD/NADH Assay Kit (MAK468; Sigma-Aldrich/Merck), which is based on an enzymatic cycling reaction coupled to reduction of MTT and colorimetric detection at 565 nm. NTC IR, NTC IR#2, and dENL/AF9 IR flies were analyzed. Because NAD and NADH require separate extraction procedures, two independent sets of 30 pooled D40 male fly heads were prepared for each biological replicate. One set was homogenized in the supplied NAD extraction buffer and the second set in NADH extraction buffer. Extracts were incubated at 60 °C for 5 min and subsequently neutralized by addition of assay buffer and the corresponding opposite extraction buffer. Samples were briefly mixed and centrifuged at 14,000 × *g* for 5 min at room temperature, and the clarified supernatants were used for analysis. A NAD standard curve ranging from 0 to 10 µM was prepared using the supplied NAD standard. Samples and standards were transferred to clear 96-well plates, followed by addition of the enzyme-based working reagent containing lactate and MTT. Absorbance at 565 nm was measured immediately after reagent addition (OD₀) and again following a 15-min incubation at room temperature (OD₁₅). The change in absorbance (ΔOD₅₆₅) was calculated for each sample and standard, and NAD and NADH concentrations were derived from the blank-corrected standard curve. The NAD/NADH ratio was calculated from four independently measured NAD and NADH concentrations. The standard curve used for quantification is shown in Supplementary Fig. 8.

### Multi-omics integration

#### Data preprocessing for matched lipidome–metabolome integration

Feature-level integration was restricted to the lipidomic and metabolomic datasets because these measurements were obtained from the same biological specimens. The analysis therefore included all matched D40 NTC IR and dENL/AF9 IR samples used for untargeted lipidomic and metabolomic profiling (n = 5 biological replicates per genotype). Each biological replicate consisted of 50 pooled fly heads.

Prior to integration, lipidomic and metabolomic features containing more than two missing or zero values across samples or having a mean intensity below 10 were removed. Duplicate annotations were collapsed by retaining the feature with the highest mean abundance. Feature intensities were log₂-transformed, zero-variance variables were removed, and variables within each omics block were scaled to unit variance before multivariate integration. These preprocessing steps were applied independently to the lipidomic and metabolomic blocks while preserving the one-to-one correspondence of matched biological samples.

#### DIABLO-based integration of matched lipidomic and metabolomic data

Matched lipidomic and metabolomic datasets were integrated using Data Integration Analysis for Biomarker discovery using Latent cOmponents (DIABLO), implemented as block sparse partial least-squares discriminant analysis (block sPLS-DA) in the mixOmics package in R v4.5.1. DIABLO was used to identify subsets of lipid and metabolite features that jointly contributed to discrimination between NTC IR and *dENL/AF9* IR samples while capturing coordinated variation between the two molecular blocks.

Model parameters, including the number of latent components and the number of variables retained from each block, were optimized by repeated cross-validation using the balanced error rate as the classification-performance metric. The final fitted model comprised two latent components. Block-specific sample projections were visualized using DIABLO individual plots, and agreement between the lipidomic and metabolomic representations of each matched sample was examined using an arrow plot. Associations between corresponding latent components from the two blocks were further assessed to characterize concordance between the lipidomic and metabolomic representations within the fitted model.

Features retained by sparse variable selection were visualized using a clustered image map (CIM) to examine genotype-associated abundance patterns across the matched samples. Cross-block relationships among DIABLO-selected lipid and metabolite features were additionally visualized using a Circos plot. Strong model-derived cross-block associations with an absolute correlation coefficient |r| ≥ 0.85 were retained for visualization. The Circos representation contained 120 selected features and 55 cross-block associations meeting this threshold.

#### Correlation-network analysis of DIABLO-selected features

To further characterize feature-level relationships underlying the integrated lipidome–metabolome signature, pairwise Pearson correlations were calculated between DIABLO-selected lipid and metabolite features across the matched biological samples. Cross-block feature pairs with |Pearson r| ≥ 0.70 were retained for construction of the lipid–metabolite correlation network. Nodes represented individual lipid or metabolite features, with node shape indicating the omics layer and node size reflecting network degree. Node color represented the log₂ fold change between dENL/AF9 IR and NTC IR samples, whereas edge color and thickness indicated the direction and magnitude of the corresponding correlation, respectively.

The resulting network was used to examine the broader organization of genotype-associated lipid–metabolite relationships rather than to infer causal or directional biochemical interactions. Particular attention was given in subsequent interpretation to metabolite features that were also identified independently in the single-metabolomics analysis, including the putatively annotated citrate feature.

### Statistical analysis

Statistical analyses of biochemical assays and selected individual molecular features were performed using GraphPad Prism version 10 (GraphPad Software, Boston, MA, USA). For comparisons between two independent groups, an unpaired two-tailed *t*-test with Welch’s correction was used. For experiments involving more than two groups or multiple experimental factors, analysis of variance (ANOVA) was applied as appropriate to the experimental design, followed by a multiple-comparisons test.

A two-sided *P* value <0.05 was considered statistically significant unless otherwise specified. The exact *P* values/significance definitions are provided in individual legends. Data are presented as mean ± standard error of the mean (SEM) unless otherwise indicated in the corresponding figure legend.

For transcriptomic analysis, multiple-testing correction was performed using the Benjamini–Hochberg procedure, with adjusted *P* <0.05 used to define differentially expressed genes. Lipidomic, metabolomic, and proteomic multivariate analyses and feature-selection thresholds were applied as described in their sections. Metabolomic pathway enrichment statistics, including raw *P* values, Holm-adjusted *P* values, FDR values, and pathway-impact scores, were retained and reported in Table 1. The criterion of −log10(*P*) >0.5 or pathway impact >0.1 was used only to highlight pathways graphically and was not considered a formal statistical-significance threshold.

## Results

### Neuronal *dENL/AF9* suppression elicits a stronger transcriptomic response in aged than young fly heads

To investigate whether neuronal *dENL/AF9* suppression is associated with age-dependent differences in gene expression, we performed RNA sequencing of heads from young (D3) and aged (D40) male flies expressing a *dENL/AF9*-targeting RNAi construct under the pan-neuronal *nSyb-GAL4* driver. Age-matched flies expressing, under the same driver, a non-targeting long double-stranded RNA directed against *luciferase* were used as controls (NTC IR), as previously validated in our characterization of neuronal *dENL/AF9* knockdown [16] (Fig. 1A). A total of 16,885 genes were detected across the transcriptomic dataset (Supplementary Table S1).

**Figure 1.**
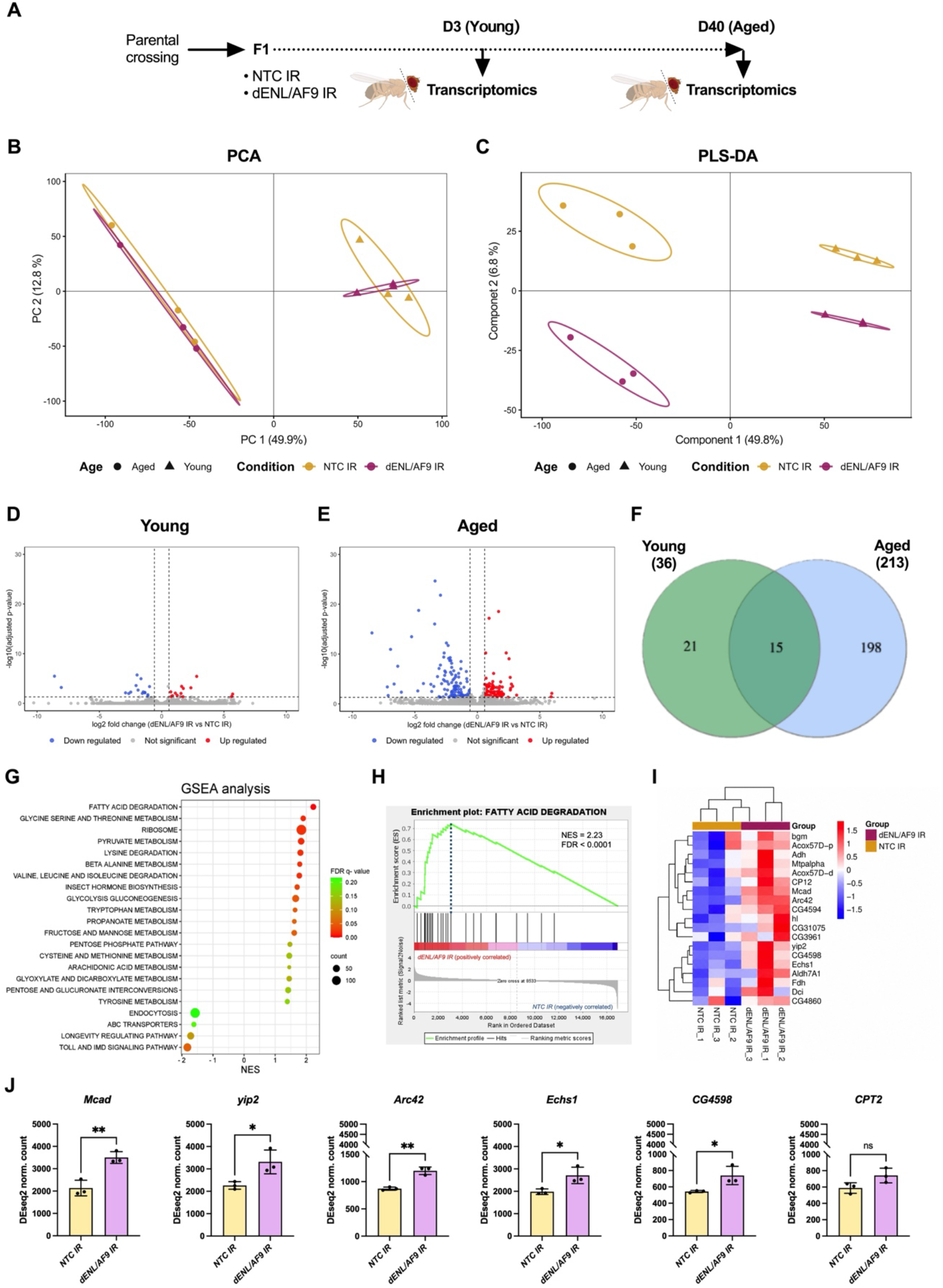
Pan-neuronal *dENL/AF9* knockdown elicits a stronger transcriptomic response in aged than young fly heads, with prominent enrichment of fatty acid degradation. (A) Experimental design for RNA sequencing (RNA-seq) of fly heads. First-generation (F1) progeny expressing either *NTC IR* or *dENL/AF9 IR* under the pan-neuronal control *nSyb-GAL4* were generated by genetic crossing and collected at D3 (Young) and D40 (Aged), corresponding to 3 and 40 days after adult eclosion, respectively. (B) and (C) Principal component analysis (PCA) and partial least-squares discriminant analysis (PLS-DA) of transcriptomic profiles from young and aged fly heads. PLS-DA model performance was assessed by cross-validation; for the two-component model, R² = 0.64 and Q² = 0.27. Each point represents one biological replicate; colours indicate experimental groups, and symbols indicate age. n = 3 biological replicates per group, with 50 pooled heads per biological replicate. (D) and (E) Volcano plots of differentially expressed genes (DEGs) in D3 and D40 *dENL/AF9*-knockdown fly heads relative to age-matched *NTC IR* controls. The x-axis shows log2 fold change (log2FC), and the y-axis shows −log10-transformed adjusted p-value (adj p-val). Red dots indicate upregulated genes, blue dots indicate downregulated genes, and grey dots indicate genes that did not meet the significance threshold (IFCI ≥ 1.5; adj p-val < 0.05). (F) Venn diagram showing the overlap of DEGs between D3 and D40 *dENL/AF9*-knockdown fly heads. (G) Gene set enrichment analysis (GSEA) of differentially regulated pathways in D40 *dENL/AF9*-knockdown fly heads. The x-axis shows the normalized enrichment score (NES), dot size represents the number of genes contributing to each pathway, and dot color indicates the false discovery rate (FDR) q-value. (H) GSEA enrichment plot for the fatty acid degradation pathway. The running enrichment score is shown across ranked genes. (I) Hierarchical clustering heatmap of leading-edge subsets from the leading-edge genes contributing to fatty acid degradation pathway enrichment. Rows represent leading-edge genes, columns represent biological replicates, and color intensity indicates scaled gene expression. (J) Normalized expression of selected fatty acid degradation-associated genes in D40 fly heads. Comparisons were performed using unpaired t-tests. Data are shown as mean ± standard deviation (SD). Statistical significance is indicated as * p<0.05,** p<0.01, and ns = not significant.

Principal component analysis (PCA) showed pronounced separation between D3 and D40 samples, indicating that age accounted for a major component of overall transcriptomic variation (Fig. 1B). Partial least-squares discriminant analysis (PLS-DA) separated the four age–genotype groups (Fig. 1C). Within each age group, genotype-associated differences between NTC IR and dENL/AF9 IR samples were visually more apparent at D40. Cross-validation of the two-component PLS-DA model yielded R² = 0.64 and Q² = 0.27, indicating modest predictive performance and supporting cautious interpretation of the supervised group separation (Supplementary Fig. 1A).

Consistent with the multivariate patterns, differential expression analysis identified substantially more differentially expressed genes (DEGs) in aged than in young *dENL/AF9*-knockdown heads. At D3, 36 genes met the predefined criteria of |fold change| ≥1.5 and adjusted *P* <0.05, compared with 213 genes at D40 (Fig. 1D, E). 21 were unique to D3 and 198 were unique to D40, whereas only 15 DEGs were shared between the two ages (Fig. 1F and Supplementary Fig. 2). Thus, the transcriptional consequences of neuronal *dENL/AF9* suppression were substantially more pronounced in aged than in young fly heads.

Gene set enrichment analysis (GSEA) further revealed distinct pathway-level patterns between the two ages. At D3, negatively enriched gene sets in the dENL/AF9 IR group were dominated by ribosome-associated processes, particularly ribosome biogenesis, together with pathways related to DNA replication and repair (Supplementary Fig. 3 and Supplementary Table S2). This ribosome-associated signature is notable because depletion of another YEATS-domain protein, YEATS2, has been shown to reduce expression of ribosomal genes in both *Drosophila* neurons and human cells [11, 31]. However, the present finding reflects pathway-level negative enrichment rather than direct evidence that the same ribosomal genes are individually downregulated following *dENL/AF9* suppression. It therefore indicates convergence on ribosome-associated transcriptional programs rather than an equivalent gene-level response. In contrast, the D40 transcriptome showed positive enrichment of multiple metabolic pathways, including fatty acid degradation, pyruvate metabolism, glycolysis/gluconeogenesis, glycine/serine/threonine metabolism, lysine degradation, branched-chain amino acid degradation, and other intermediary metabolic pathways (Fig. 1G). Notably, the KEGG ribosome pathway was also positively enriched at D40, contrasting with the negative ribosome-associated enrichment observed at D3 and further indicating that the pathway-level transcriptional response to *dENL/AF9* suppression differed between young and aged animals.

Among the D40 metabolic signatures, fatty acid degradation showed particularly strong positive enrichment (NES = 2.23, FDR *q* <0.0001; Fig. 1G, H). Leading-edge analysis identified a coordinated subset of genes contributing to this enrichment (Fig. 1I). Consistent with the pathway-level result, representative fatty acid degradation- and β-oxidation-associated genes, including *Mcad*, *yip2*, *Arc42*, *Echs1*, and *CG4598*, showed higher normalized expression in aged *dENL/AF9*-knockdown heads than in age-matched NTC IR controls, whereas the increase in *CPT2* did not reach statistical significance (Fig. 1J). These findings identify fatty acid degradation-associated transcription as one of the prominent features of the aged *dENL/AF9*-knockdown transcriptome.

GSEA also identified negative enrichment of the KEGG longevity-regulating pathway at D40 (NES = −1.72, FDR *q* = 0.10; Fig. 1G and Supplementary Fig. 4A, B), representing an exploratory enrichment signal at the predefined GSEA threshold. Although this KEGG pathway contains components of several established longevity pathways, including insulin/insulin-like signalling (IIS), examination of the leading-edge subset showed that the negative enrichment was driven predominantly by reduced expression of heat-shock-related genes, particularly *Hsp70Ba*, *Hsp70Bb*, and *Hsp70Bbb* (Supplementary Fig. 4C, D). Canonical IIS components were not prominent among the leading-edge genes. This observation is consistent with our previous study, in which neuronal *dENL/AF9* suppression did not significantly reduce representative canonical IIS components at D3 or D20 [16], and provides no evidence from the current D40 transcriptome for coordinated suppression of canonical IIS signalling.

Together, these analyses show that neuronal *dENL/AF9* suppression is associated with a substantially stronger transcriptional response in aged than in young fly heads. At D40, this response is distinguished by positive enrichment of multiple intermediary metabolic programs, with fatty acid degradation representing one of the strongest signatures, whereas the exploratory longevity-pathway signal is driven primarily by Hsp70-family genes rather than canonical IIS components.

### Aged *dENL/AF9*-knockdown heads exhibit a distinct lipidomic profile

Given the enrichment of fatty-acid-degradation-associated transcriptional programs in aged *dENL/AF9*-knockdown heads, together with our previous observation that neuronal *dENL/AF9* suppression reduces lipid peroxidation [16], we next examined whether this phenotype was accompanied by alterations in the aged head lipidome.

Untargeted lipidomics was performed on matched D40 head samples from NTC IR and dENL/AF9 IR flies (Fig. 2A). After quality-control filtering, lipidomic profiling retained 5,583 features, comprising 3,692 features detected in positive-ion mode and 1,891 in negative-ion mode.

**Figure 2.**
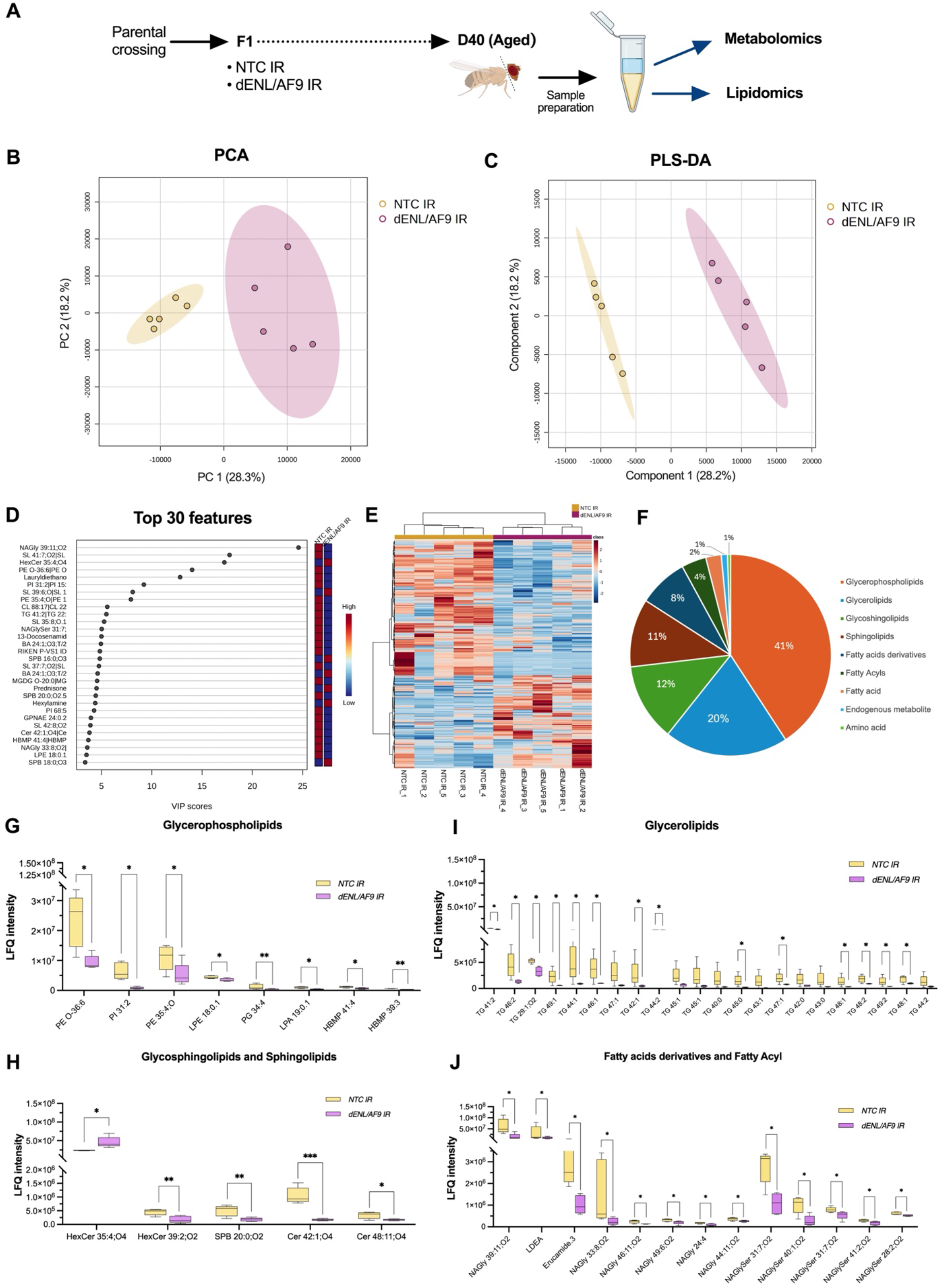
Aged *dENL/AF9*-knockdown fly heads exhibit a distinct lipidomic profile with class- and species-specific abundance differences. **(A)** Experimental design for untargeted lipidomic and metabolomic profiling of aged fly heads. Heads from D40 F1 flies expressing either NTC IR or dENL/AF9 IR were collected and processed for LC–MS-based lipidomic and metabolomic analyses. **(B)** and **(C)** Principal component analysis (PCA) and partial least-squares discriminant analysis (PLS-DA), respectively, of lipidomic profiles from aged NTC IR and dENL/AF9 IR fly heads. Each point represents one biological replicate; n = 5 biological replicates per group, with 50 pooled heads per biological replicate. PLS-DA model performance was assessed by cross-validation; for the two-component model, R² = 0.99 and Q² = 0.79. **(D)** Variable importance in projection (VIP) plot showing the top 30 lipid features contributing to group discrimination in the PLS-DA model. The adjacent heat strip indicates relative feature abundance between groups. **(E)** Hierarchical clustering heatmap of lipid features with VIP scores > 1 in aged NTC IR and dENL/AF9 IR fly heads. **(F)** Pie chart showing the distribution of annotated lipid features with VIP scores > 1 according to lipid category. **(G-J)** Relative abundance of representative altered lipid species classified as glycerophospholipids **(G)**, glycosphingolipids and sphingolipids **(H)**, glycerolipids (I), and fatty acid derivatives and fatty acyls (J). Lipid identities represent database-based annotations and were not independently confirmed using authentic reference standards. Comparisons were performed using multiple unpaired t-tests with Bonferroni correction. Statistical significance is indicated as * p<0.05,** p<0.01, and *** p<0.001.

PCA indicated genotype-associated differences in the overall lipidomic profiles, whereas PLS-DA more clearly separated NTC IR and dENL/AF9 IR samples (Fig. 2B,C). Cross-validation across increasing numbers of latent components selected a two-component model as optimal (Supplementary Fig. 1B). VIP-based feature selection further highlighted the lipid features contributing most strongly to genotype discrimination. Figure 2D displays the top 30 VIP-ranked lipid features, whereas 285 features across the complete dataset had VIP scores >1 and were retained for subsequent analysis (Supplementary Table S3). Hierarchical clustering of these 285 VIP features revealed a highly distinct and contrastive abundance pattern between the two genotypes (Fig. 2E). NTC IR and dENL/AF9 IR replicates formed clearly separated clusters in the column dendrogram, while coordinated groups of lipid features displayed opposing relative-abundance patterns between the genotypes. These results indicate that genotype-associated variation was a major source of structure among the VIP-selected features and define a clear lipidomic signature distinguishing aged *dENL/AF9*-knockdown heads from control group.

The 285 total VIP features, of which 201 could be assigned to the annotation categories presented in Suplementary Table S3. They were distributed across multiple lipid categories, with glycerophospholipids and glycerolipids representing the largest groups, together with glycosphingolipids, sphingolipids, fatty acid derivatives, fatty acyls, and other lipid-associated features (Fig. 2F). To examine whether genotype-associated abundance patterns differed among lipid classes, the annotated VIP features were further stratified according to lipid category and examined by hierarchical clustering (Supplementary Fig. 5A–F). These class-specific analyses revealed distinct and heterogeneous abundance patterns between NTC IR and dENL/AF9 IR samples across the major lipid categories. For visualization of individual features, up to 15 VIP lipid features within each category were displayed alongside the corresponding heatmaps (Supplementary Fig. 5A–F), and features showing significant between-genotype abundance differences were selected for presentation in the main figure (Fig. 2G–J).

Among glycerophospholipids, significantly altered features included species showing both lower and higher abundance in dENL/AF9 IR heads, consistent with a heterogeneous class-level response (Fig. 2G and Supplementary Fig. 5A). Glycosphingolipid and sphingolipid features similarly exhibited mixed genotype-associated abundance differences (Fig. 2H and Supplementary Fig. 5B, C). In contrast, the glycerolipid analysis revealed a conspicuous cluster of triglyceride (TG) features with consistently lower abundance in dENL/AF9 IR samples (red dashed box, Supplementary Fig. 5D). These clustered TG species, together with additional TG features showing significant genotype-associated differences, were selected for display in Fig. 2I. Several fatty acid derivative and fatty acyl features also showed significantly lower abundance in *dENL/AF9*-knockdown heads and were highlighted in Fig. 2J (Supplementary Fig. 5E, F). Together, these analyses show that aged *dENL/AF9*-knockdown heads exhibit a distinct lipidomic profile characterized by class- and species-specific abundance differences rather than a uniform shift across the lipidome. A particularly prominent feature was the lower abundance of multiple TG-related species, accompanied by reductions in several fatty acid derivative and fatty acyl features, whereas glycerophospholipid- and sphingolipid-associated features showed more heterogeneous patterns.

### Aged *dENL/AF9*-knockdown heads exhibit a distinct metabolomic profile with features mapping to central carbon, nicotinamide, and purine metabolism

Positive enrichment of multiple metabolic pathways in the aged transcriptomic dataset, together with the distinct lipidomic profile observed at D40, prompted us to examine whether neuronal *dENL/AF9* suppression was also associated with broader changes in small-molecule metabolism. We therefore analyzed the untargeted metabolomic profiles obtained from the same D40 head samples used for lipidomic analysis (Fig. 2A). Following quality-control filtering, 1,359 metabolite features were retained, comprising 937 features detected in positive-ion mode and 422 in negative-ion mode. PCA revealed overlapping but genotype-associated variation in the aged metabolomic dataset (Fig. 3A). The PLS-DA score plot based on the first two latent components showed clearer genotype-associated separation (Fig. 3B). Cross-validation identified four components as optimal (Supplementary Fig. 1C); the two-component model shown in Fig. 3B had R² = 0.97 and Q² = 0.59. Figure 3C shows the top 30 VIP-ranked features contributing to genotype discrimination, while 198 features across the dataset had VIP scores >1 and were retained for subsequent analysis. Hierarchical clustering of these 198 VIP features revealed distinct abundance patterns between NTC IR and dENL/AF9 IR samples (Fig. 3D), further defining a genotype-associated metabolomic signature in aged fly heads.

**Figure 3.**
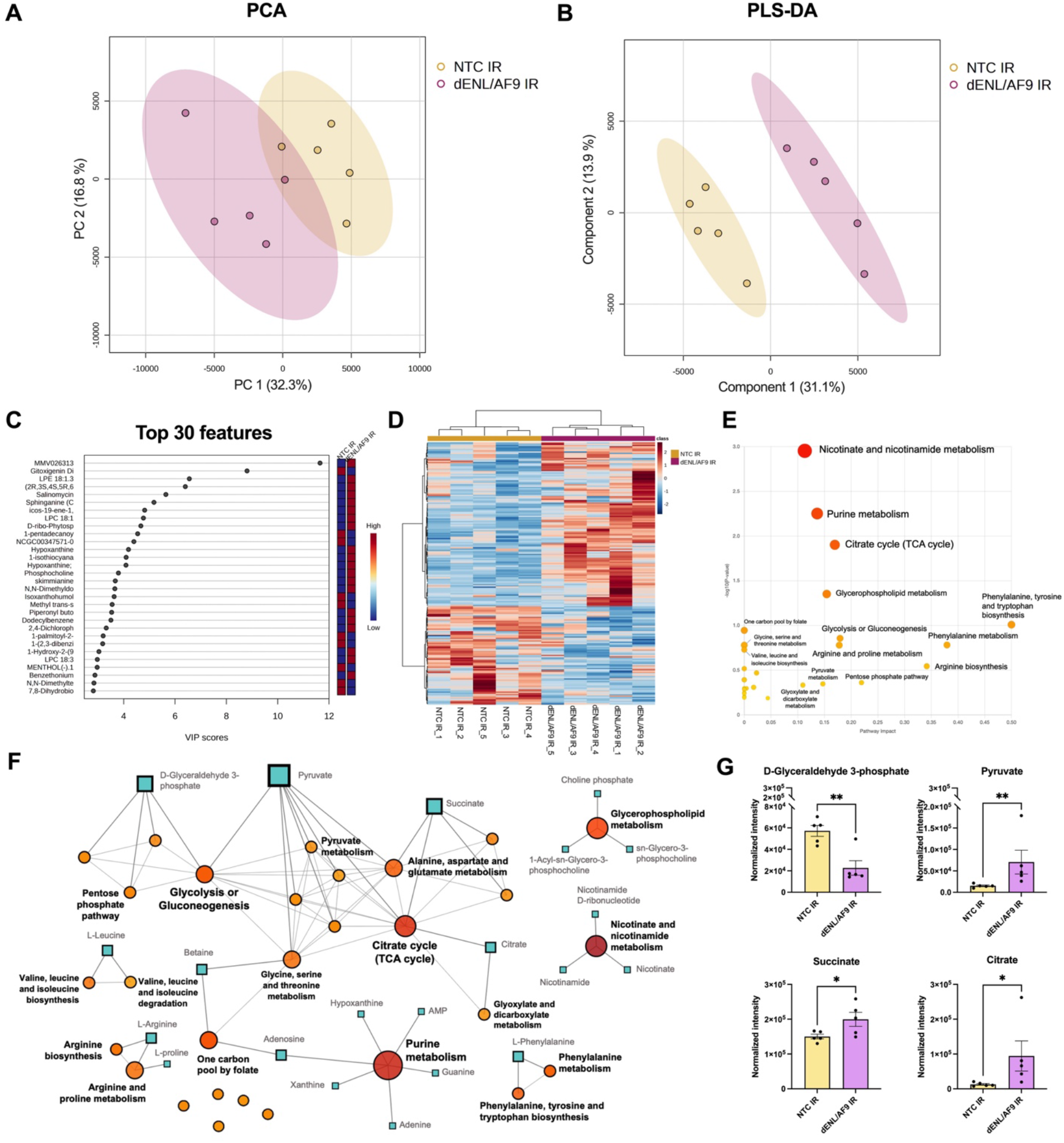
Aged *dENL/AF9*-knockdown fly heads exhibit a distinct metabolomic profile with features mapping to central carbon, nicotinamide, and purine metabolism. **(A)** and **(B)** Principal component analysis (PCA) and partial least-squares discriminant analysis (PLS-DA), respectively, of untargeted metabolomic profiles from aged NTC IR and dENL/AF9 IR fly heads. Each point represents one biological replicate. n = 5 biological replicates per group, with 50 pooled heads per biological replicate. **(C)** Variable importance in projection (VIP) plot showing the top 30 metabolite features contributing to group discrimination in the PLS-DA model. The adjacent heat strip indicates relative feature abundance between groups. PLS-DA model performance was assessed by cross-validation; for the two-component model, R² = 0.97 and Q² = 0.59. (D) Hierarchical clustering heatmap of metabolite features with VIP scores > 1 in aged NTC IR and dENL/AF9 IR fly heads. (E) Pathway analysis of important metabolite features. Each pathway is represented by a circle positioned according to −log10(P value) on the vertical axis and pathway impact on the horizontal axis. Circle color represents the statistical significance of pathway enrichment, whereas circle size reflects pathway impact. Highlighted pathways met either −log10(P value) > 0.5 or pathway impact > 0.1. **(F)** Metabolite–pathway interaction network showing relationships between altered metabolite features and associated metabolic pathways. Circles represent metabolic pathways, squares represent individual metabolites, and edges indicate known biochemical relationships. Node size is proportional to the number of connected edges. **(G)** Relative abundance of metabolites associated with central carbon metabolism. Metabolite identities represent putative annotations based on the LC–MS/MS database-matching workflow and were not confirmed using authentic reference standards. Comparisons were performed using unpaired t-tests. Data are shown as mean ± standard error of the mean (SEM). Statistical significance is indicated as * p<0.05 and ** p<0.01.

To identify the metabolic processes represented within this signature, all 198 VIP features were submitted to pathway analysis using the *D. melanogaster* KEGG library in MetaboAnalyst. Of these, 80 features could be mapped to KEGG compounds and therefore contributed to the pathway analysis; their annotations and relative abundances are reported in Supplementary Table S4. The resulting analysis highlighted several metabolic processes (Fig. 3E and Table 1). The strongest nominal enrichment was observed for nicotinate and nicotinamide metabolism, followed by purine metabolism and the citrate cycle (TCA cycle), with additional signals involving glycerophospholipid metabolism and several amino acid- and central-carbon-associated pathways.

The metabolite–pathway interaction network further showed that these pathway signals were organized into several interconnected metabolic modules (Fig. 3F). A prominent cluster involved central carbon metabolism, linking glycolysis/gluconeo-genesis, pyruvate metabolism, the TCA cycle, alanine, aspartate, and glutamate metabolism, glyoxylate and dicarboxylate metabolism, and the pentose phosphate pathway. To represent this highly interconnected cluster, four putatively annotated metabolites—D-glyceraldehyde-3-phosphate, pyruvate, succinate, and citrate—were selected for individual visualization in Fig. 3G. Pyruvate, citrate, and succinate features showed higher abundance in dENL/AF9 IR heads, whereas the D-glyceraldehyde-3-phosphate-associated feature showed lower abundance (Fig. 3G and Supplementary Table S4).

Relative abundances of VIP metabolites contributing to other pathway signals are shown in Supplementary Fig. 6. Multiple purine-associated features differed between the two genotypes, including metabolites putatively annotated as adenine, guanine, xanthine, hypoxanthine, AMP, and adenosine (Supplementary Fig. 6 and Supplementary Table S4). Features mapping to nicotinate and nicotinamide metabolism included putative nicotinate, nicotinamide, and β-nicotinamide mononucleotide, which also showed genotype-associated abundance differences between NTC IR and dENL/AF9 IR heads.

Following the primary pathway analysis, we additionally examined the metabolomic dataset for metabolites biologically related to the pathways emerging from the 80 KEGG-mapped VIP features. This pathway-guided inspection identified six additional features annotated as tryptophan, alanine, fumarate, 5-hydroxyindole-3-acetic acid, N-acetylaspartate (NAA), and inosine-5′-monophosphate (IMP) (Supplementary Fig. 6 and Supplementary Table S4). These additional features were examined to provide further context for the pathway-level findings and were not included among the 80 mapped metabolites used to calculate the pathway-enrichment statistics. Among these additional observations, the putatively annotated NAA, which is linked to neuronal carbon metabolism in mammals [32], showed higher abundance in *dENL/AF9*-knockdown heads. Together, these analyses identify a distinct genotype-associated metabolomic profile in aged *dENL/AF9*-knockdown heads, characterized by features mapping to central carbon, nicotinamide, purine, and amino acid metabolism.

### Proteomic and biochemical analyses identify an energetic and redox signature in aged *dENL/AF9*-knockdown heads

Transcriptomic, lipidomic, and metabolomic analyses identified a genotype-associated metabolic signature in aged dENL/AF9-knockdown heads, characterized by enrichment of fatty acid degradation and other metabolism-associated transcriptional programs, class- and species-specific differences in lipid abundance, and changes in putatively annotated metabolites mapping to central carbon, nicotinamide, purine, and amino acid metabolism. To determine whether this metabolic signature was also represented at the protein level, we next performed untargeted proteomic profiling of independent D40 NTC IR and dENL/AF9 IR head samples (Fig. 4A). A total of 287 proteins were retained for downstream analysis.

**Figure 4.**
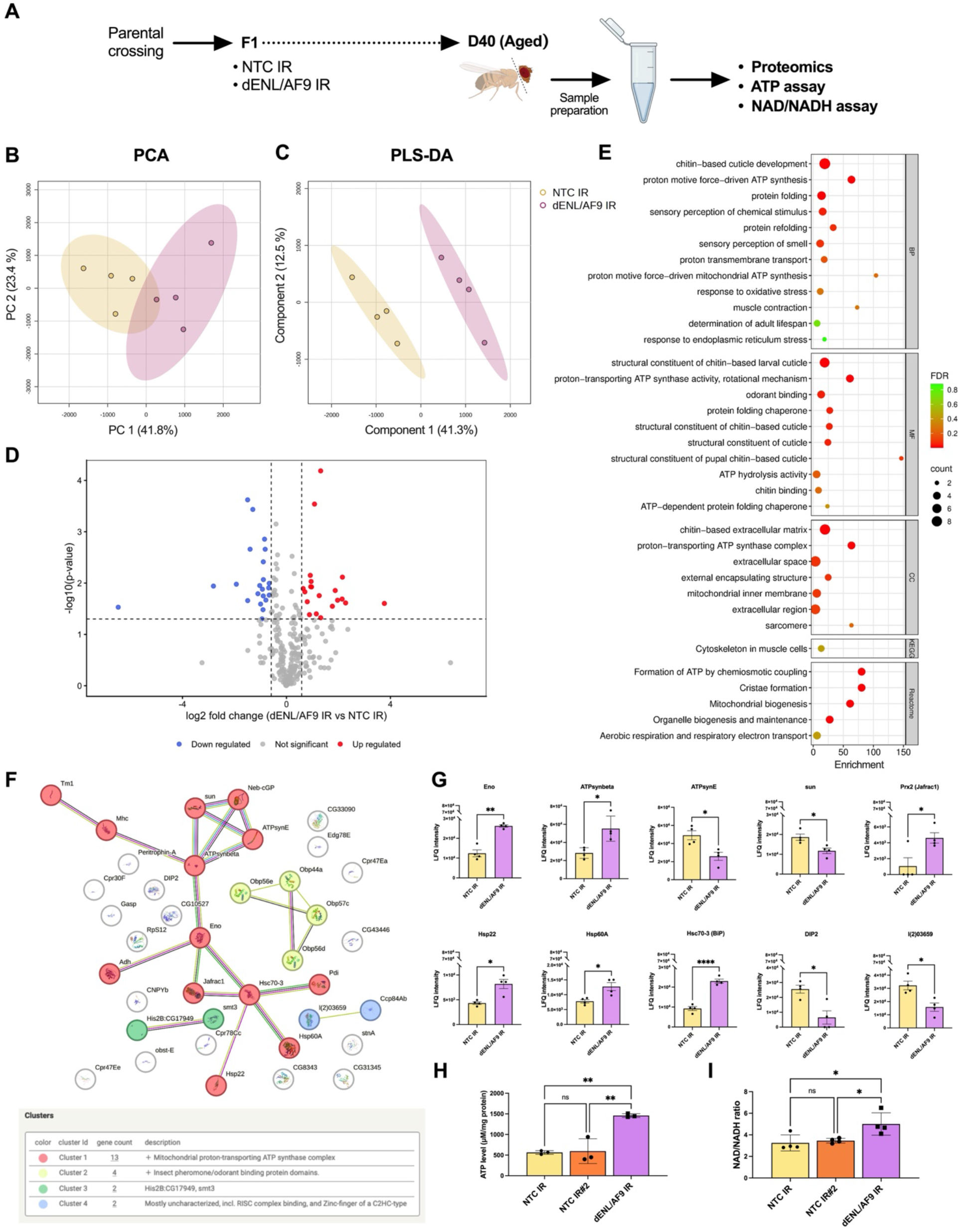
Proteomic and biochemical analyses identify an energetic and redox signature in aged *dENL/AF9*-knockdown fly heads. **(A)** Experimental overview of aged fly head collection for proteomic profiling, ATP measurement, and nicotinamide adenine dinucleotide/nicotinamide adenine dinucleotide hydride (NAD/NADH) ratio assessment. **(B)** and **(C)** Principal component analysis (PCA) and partial least-squares discriminant analysis (PLS-DA), respectively, of proteomic profiles from aged NTC IR and dENL/AF9 IR fly heads. Each point represents one biological replicate. n = 4 biological replicates per group, with 50 pooled heads per biological replicate. PLS-DA model performance was assessed by cross-validation; for the two-component model, R² = 0.98 and Q² = 0.74. **(D)** Volcano plot of differentially abundant proteins (DAPs) in D40 *dENL/AF9*-knockdown fly heads relative to age-matched NTC IR controls. The x-axis shows log2 fold change (log2FC), and the y-axis shows −log10-transformed P value. Red and blue dots indicate significantly increased and decreased proteins, respectively; gray dots indicate proteins that did not meet the significance criteria (IFCI ≥ 1.5; p-val < 0.05). **(E)** Functional enrichment analysis of DAPs using DAVID. Enriched terms are derived from Gene Ontology Biological Process (BP), Molecular Function (MF), and Cellular Component (CC), together with Kyoto Encyclopedia of Genes and Genomes (KEGG) and Reactome annotations. The x-axis shows the enrichment score, dot size represents the number of proteins assigned to each term, and dot color indicates the FDR q value. **(F)** Protein– protein interaction (PPI) network of DAPs generated using the STRING database. Nodes represent proteins and edges indicate known or predicted protein associations. Functionally related proteins are organized into clusters, with major clusters summarized in the accompanying table. **(G)** Relative abundance of selected differentially abundant proteins highlighted by the PPI network and functional-enrichment analyses, including proteins associated with mitochondrial ATP synthesis, energy metabolism, and cellular stress/protein-folding processes. Relative abundance values represent normalized protein intensities and are shown as mean ± SEM. Comparisons were performed using unpaired t-tests with Welch’s correction. **(H)** and **(I)** ATP content and NAD/NADH ratio, respectively, in aged fly heads. NTC IR#2 denotes an independent *mCherry* RNAi control expressed under the *nSyb* driver. Holm-Šídák’s multiple comparisons test was used for the comparisons. Data are presented as mean ± SD. Statistical significance is indicated as * p<0.05 and ** p<0.01.

PCA showed overlapping proteomic distributions with some genotype-associated variation, while the PLS-DA score plot based on the first two latent components showed clearer separation between NTC IR and dENL/AF9 IR samples (Fig. 4B, C). Cross-validation identified four components as optimal (Supplementary Fig. 1D); the two-component model shown in Fig. 4C had R² = 0.98 and Q² = 0.75. Differential abundance analysis identified 42 differentially abundant proteins (DAPs) in aged *dENL/AF9*-knockdown heads relative to controls, including both increased and decreased proteins (Fig. 4D and Supplementary Table S5).

Functional enrichment analysis of the DAPs identified multiple biological terms related to mitochondrial energy metabolism and ATP-associated processes, including proton-motive-force-driven ATP synthesis, electron transport, mitochondrial protein-containing complexes, and ATP hydrolysis (Fig. 4E). Notably, the GO biological-process term “determination of adult lifespan” was also enriched and included Hsp22, Prx2, and stunted (sun), three proteins previously implicated in the regulation of Drosophila longevity [33–35]. Protein–protein interaction (PPI) analysis further organized the DAPs into functionally connected clusters, including a prominent cluster containing proteins associated with mitochondrial energy metabolism and ATP synthesis (Fig. 4F). Examination of individual DAPs represented within the PPI network and enriched functional categories showed significant abundance changes among selected ATP synthase-associated components (Fig. 4G). The relative abundances of additional DAPs assigned to the major STRING cluster are shown in Supplementary Fig. 7. Together, the enrichment of mitochondrial and ATP-associated functional categories and the altered abundance of multiple proteins within the corresponding interaction network indicate that the metabolic signature identified by transcriptomic, lipidomic, and metabolomic profiling is also represented at the protein level in aged *dENL/AF9*-knockdown heads, with prominent involvement of mitochondrial energy metabolism and ATP-associated processes.

To independently assess this energy-associated signature at the biochemical level, we quantified ATP abundance in a separate cohort of D40 fly heads. We first compared *dENL/AF9* RNAi flies with the *luciferase*-directed RNAi control used throughout the transcriptomic, lipidomic, metabolomic, and proteomic analyses (NTC IR); both RNAi constructs were expressed under the pan-neuronal *nSyb*-GAL4 driver. To determine whether the ATP phenotype was reproducible relative to an additional RNAi control construct, we also included age-matched flies expressing *mCherry*-directed RNAi (NTC IR#2). ATP abundance was comparable between the two control groups but was significantly higher in *dENL/AF9*-knockdown heads than in either control group (Fig. 4H), providing biochemical support for the genotype-associated energy-related signature identified by the molecular profiling analyses.

Because the metabolomic analysis also identified features mapping to central-carbon and nicotinate/nicotinamide metabolism, and because NAD(H) couples intermediary metabolism to cellular redox balance and mitochondrial electron transfer, we next assessed the NAD/NADH ratio in independent D40 cohorts using the same three genotypes. The NAD/NADH ratio was comparable between NTC IR and NTC IR#2 but was significantly higher in *dENL/AF9* IR heads relative to both control groups (Fig. 4I), consistent with a shift toward a more oxidized whole-head NAD redox state. Calibration curves used for ATP and NAD quantification are shown in Supplementary Fig. 8.

Together, the D40 datasets define a genotype-associated metabolic and energetic signature, accompanied by an altered NAD redox state in *dENL/AF9*-knockdown fly heads. Transcriptomic profiling identified enrichment of fatty acid degradation- and other metabolism-associated gene programs; lipidomics revealed class- and species-specific differences, including lower abundance of multiple TG-related, fatty acid derivative, and fatty acyl features; metabolomics identified features mapping to central carbon, nicotinamide, purine, and amino acid metabolism; and proteomics implicated mitochondrial and ATP-associated processes. The increased ATP abundance and NAD/NADH ratio provide targeted biochemical evidence that this molecular signature is accompanied by altered whole-head energetic and redox states.

### Matched lipidome–metabolome integration identifies a joint genotype-associated feature signature

The individual lipidomic and metabolomic analyses identified distinct genotype-associated molecular patterns in aged *dENL/AF9*-knockdown heads. Because these datasets were generated from the same biological specimens, we next asked whether lipid and metabolite features formed a coordinated cross-block structure across matched samples. We therefore applied DIABLO-based block sparse partial least-squares discriminant analysis (block sPLS-DA) to the matched D40 lipidomic and metabolomic datasets from NTC IR and dENL/AF9 IR heads (Fig. 5A). This analysis was used to identify subsets of lipid and metabolite features that jointly contributed to genotype discrimination while capturing coordinated variation between the two molecular layers.

**Figure 5.**
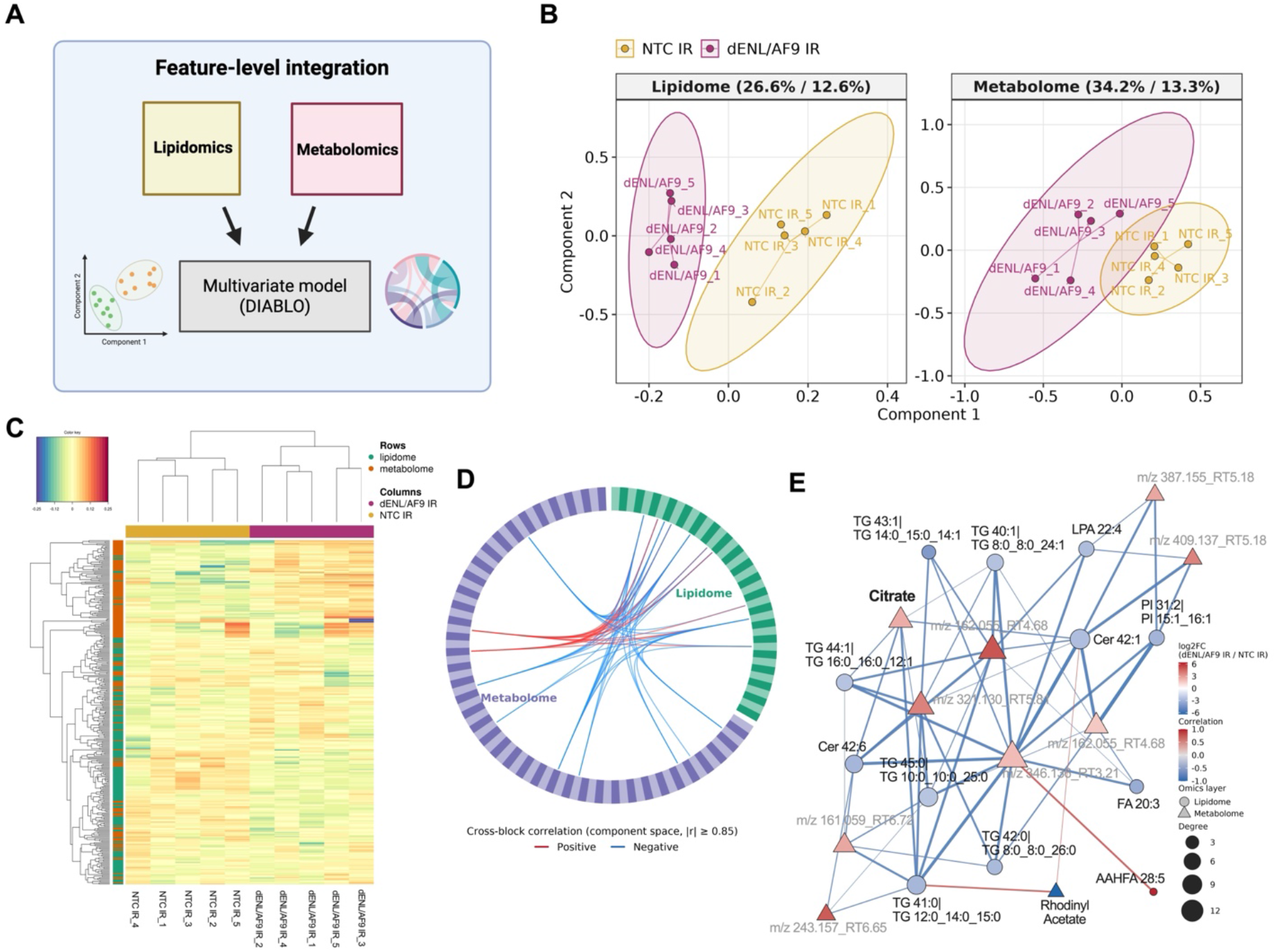
Matched lipidome–metabolome integration identifies a joint genotype-associated feature signature linking citrate with altered lipid features in aged *dENL/AF9*-knockdown fly heads. DIABLO integrated analyses were performed using matched lipidomic and metabolomic datasets from aged *NTC IR* and *dENL/AF9 IR* lines. **(A)** Schematic overview of matched lipidome–metabolome feature-level integration using DIABLO (Data Integration Analysis for Biomarker discovery) **(B)** Block-specific score plots from the DIABLO multi-block sparse partial least-squares discriminant analysis (sPLS-DA) model for aged lipidomics and metabolomics datasets derived from same biological samples. Each point represents one biological replicate comprising 50 pooled heads, and colors indicate experimental groups. Ellipses illustrate group clustering within each dataset. All five replicates per group were included from each omics block. Percentages shown in parentheses after each omics-block name indicate the proportion of variance explained by Component 1 and Component 2, respectively (lipidome: 26.6% and 12.6%; metabolome: 34.2% and 13.3%). **(C)** Clustered Image Map (CIM) generated with the cimDiablo function, showing the molecular signatures of the model’s key variables. Samples are shown in columns and selected features associated with the first component in rows. Hierarchical clustering of rows and columns was performed using Euclidean distance and complete linkage. **(D)** Circos representation of cross-block associations among DIABLO-selected lipidomic and metabolomic features. The outer ring contains 120 selected features, with lipidomic and metabolomic features shown in separate sectors. Connecting lines indicate strong cross-block associations meeting |r| ≥ 0.85; red and blue lines indicate positive and negative associations, respectively. **(E)** Cross-block correlation network of DIABLO-selected lipid and metabolite features. Nodes represent lipid or metabolite features; node shape indicates the omics layer, node color reflects log₂FC, and node size is proportional to network degree. Edge color indicates the direction of association, and edge thickness reflects correlation strength. Only associations meeting |Pearson *r*| ≥ 0.70 are displayed. Metabolite features with low-confidence annotations or without supporting MS/MS information are labelled using their *m/z* and retention time (RT) rather than tentative compound names. Named metabolites and lipids are shown using their database-based annotations; metabolite identities remain putative unless otherwise stated.

Within the fitted multiblock model, both the lipidomic and metabolomic blocks showed separation between NTC IR and dENL/AF9 IR samples along the first two latent components (Fig. 5B). In the lipidomic block, Component 1 and Component 2 accounted for 26.6% and 12.6% of the block variance, respectively, whereas the corresponding components in the metabolomic block accounted for 34.2% and 13.3%, respectively. The corresponding DIABLO arrow plot illustrated the relative projections of the two molecular blocks for each matched biological sample (Supplementary Fig. 9A). The latent components derived from the lipidomic and metabolomic blocks were positively correlated, with correlations of 0.84 for component 1 and 0.76 for component 2 (Supplementary Fig. 9B, C), indicating concordance between the two data representations within the integrated model.

Sparse feature selection identified subsets of lipid and metabolite features contributing to genotype discrimination. The clustered image map (CIM) generated from these DIABLO-selected features showed distinct genotype-associated abundance patterns and separation of NTC IR and dENL/AF9 IR samples across the joint feature set (Fig. 5C). Cross-block relationships among the selected features were further visualized using a Circos plot, which identified 55 strong positive and negative lipid–metabolite associations within the fitted model (|r| ≥ 0.85; Fig. 5D). These associations involved multiple metabolite and lipid features rather than being dominated by a single metabolite, supporting a broader coordinated genotype-associated structure across the matched molecular layers.

To further characterize the feature-level relationships underlying the cross-block associations visualized in the Circos plot, we next constructed a lipid–metabolite correlation network from the DIABLO-selected features (Fig. 5E). Low-confidence metabolite annotations, including features lacking supporting MS/MS information, are displayed in the network using their *m/z*–retention-time identifiers rather than tentative compound names to avoid overinterpretation of individual metabolite identities. Among the connected metabolite features, the putatively annotated citrate feature was of particular interest because it had also been identified independently in the metabolomic analysis as part of the central-carbon-associated signature, and its annotation was supported by a mass-consistent LC–MS precursor signal. Citrate was therefore examined as a recurrent and biologically interpretable component of the integrated signature rather than as a uniquely connected network hub. Within the matched lipid–metabolite association network, citrate showed inverse associations with several altered lipid features, including triglyceride- and ceramide-associated species (Fig. 5E and Table 2). Specifically, citrate abundance was inversely associated with TG 41:0, Cer 42:6, Cer 42:1, TG 44:1, TG 45:0, and TG 40:1, with correlation coefficients ranging from −0.70 to −0.83 (Table 2). Thus, the recurrence of citrate across the individual metabolomic and matched integrative analyses places this central-carbon-associated feature within the broader lipid–metabolite signature observed in *dENL/AF9*-knockdown heads.

**Table 2.** Lipid features associated with the putatively annotated citrate feature in the matched lipidome–metabolome integration network.

| From |  | To |  | Correlation |
| --- | --- | --- | --- | --- |
| block | feature | block | feature |  |
| metabolome | CITRATE | lipidome | TG 41:0\|TG 12:0_14:0_15:0 | -0.83 |
| metabolome | CITRATE | lipidome | Cer 42:6 | -0.79 |
| metabolome | CITRATE | lipidome | Cer 42:1 | -0.76 |
| metabolome | CITRATE | lipidome | TG 44:1\|TG 16:0_16:0_12:1 | -0.75 |
| metabolome | CITRATE | lipidome | TG 45:0\|TG 10:0_10:0_25:0 | -0.73 |
| metabolome | CITRATE | lipidome | TG 40:1\|TG 8:0_8:0_24:1 | -0.70 |

Overall, the matched lipidome–metabolome analysis indicates that genotype-associated lipidomic and metabolomic differences are organized within a shared multivariate structure across the same D40 biological samples. Citrate represents one recurrent and biologically interpretable component of this broader signature, linking the central-carbon-associated metabolomic pattern with accompanying lipid-associated changes.

## Discussion

The present study extends our previous identification of neuronal *dENL/AF9* as a regulator of lifespan, healthspan, and oxidative-stress resilience [16] by defining the molecular state associated with its suppression in aged flies. As summarized in Figure 6, independently generated molecular datasets converge on a metabolic and energetic signature characterized by enrichment of fatty-acid-degradation and other intermediary metabolic transcriptional programs, broad lipid-associated changes, alterations in central-carbon, nicotinamide, and purine-associated metabolites, remodeling of mitochondrial and ATP-related protein processes, increased ATP abundance, and an elevated NAD/NADH ratio. This convergence indicates that neuronal *dENL/AF9* suppression is associated with coordinated changes across several molecular layers rather than with an isolated transcriptional or metabolic phenotype. Importantly, only the transcriptomic dataset directly compared young and aged animals and showed a substantially stronger response at D40 than at D3. The lipidomic, metabolomic, proteomic, and biochemical datasets therefore define a genotype-associated molecular state in aged fly heads rather than metabolic remodeling across aging. Accordingly, these features should be interpreted as molecular correlates of the previously established longevity phenotype rather than as demonstrated causal mediators of lifespan extension.

**Figure 6.**
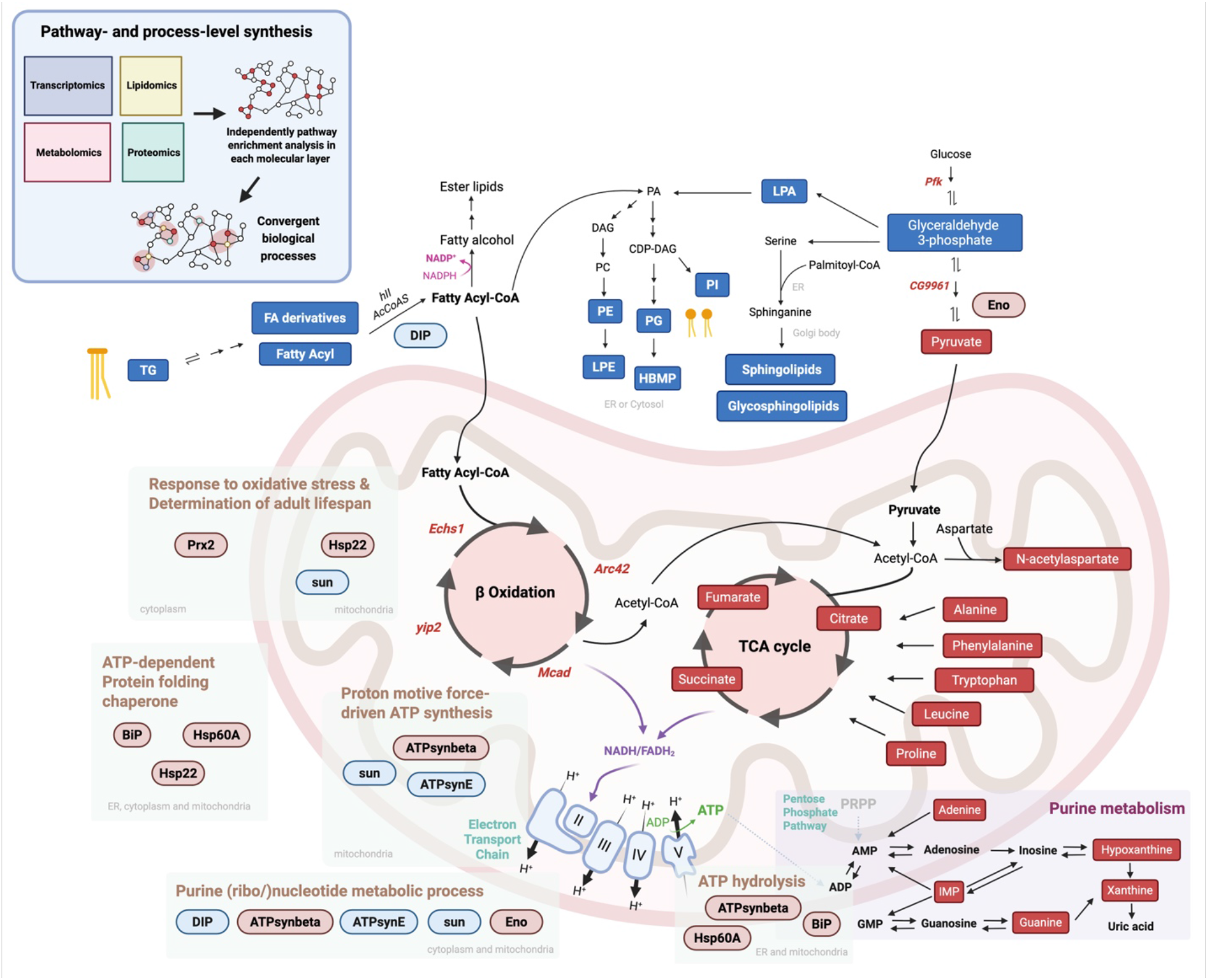
Cross-omics pathway and process synthesis defines a coherent metabolic and energetic signature in aged *dENL/AF9*-knockdown fly heads. Principal findings from the transcriptomic, lipidomic, metabolomic, proteomic, and biochemical analyses were summarized to identify convergent biological processes associated with pan-neuronal *dENL/AF9* suppression at D40. Transcriptomic profiling showed positive enrichment of fatty acid degradation and other intermediary metabolic programs, including increased expression of representative fatty acid degradation-associated genes such as *Echs1, yip2, Mcad,* and *Arc42*. Lipidomic profiling identified class- and species-specific abundance differences, including lower abundance of multiple triglyceride (TG)- related, fatty acid derivative, and fatty acyl features, together with heterogeneous changes among glycerophospholipid and sphingolipid-associated features. Metabolomic profiling identified putatively annotated features mapping to central-carbon, amino acid, nicotinamide, and purine metabolism, including pyruvate, citrate, succinate, and other pathway-associated metabolites. Proteomic analysis independently highlighted mitochondrial energy metabolism and ATP-associated processes, including proton-motive-force-driven ATP synthesis, electron transport, ATP hydrolysis, and the GO biological-process term “determination of adult lifespan.” Increased ATP abundance and an elevated NAD/NADH ratio provided supporting biochemical evidence of altered whole-head energetic and redox states. Collectively, these independently analyzed molecular layers converge on a D40 genotype-associated signature involving lipid handling, central-carbon and nucleotide metabolism, mitochondrial energy-associated processes, and redox regulation. Red labels indicate higher abundance/expression and blue labels lower abundance/expression in dENL/AF9 IR relative to NTC IR, whereas uncoloured metabolites and pathway arrows indicate biochemical relationships included for pathway context. The figure was created using ©BioRender (https://biorender.com).

The pronounced difference between the D3 and D40 transcriptomic responses suggests that the consequences of reducing this YEATS-domain histone-acylation reader depend strongly on physiological context. This interpretation is consistent with our recent study of pharmacological ENL/AF9 YEATS-domain inhibition in neuronal stress models [12]. In differentiated human neuronal cells, the selective inhibitor SR-0813 improved survival during oxidative, proteotoxic, and particularly endoplasmic-reticulum stress, but exacerbated toxicity during rotenone-induced mitochondrial dysfunction [12]. In *Drosophila* neurodegeneration models, reduction of ENL/AF9 activity was likewise beneficial in some disease contexts but showed limited or detrimental effects in others [12]. Although those experiments differ from the present study in both intervention and biological setting, they provide independent evidence that the consequences of reducing ENL/AF9 activity are highly dependent on the prevailing metabolic, redox, and stress environment. The stronger transcriptional response observed in aged *dENL/AF9*-knockdown heads is therefore consistent with a broader model in which the functional output of ENL/AF9 depends on the physiological state of the nervous system rather than reflecting control of a fixed transcriptional program.

This context dependence may also help reconcile the apparent paradox that endogenous *dENL/AF9* expression declines with age and following oxidative challenge [16], whereas further neuronal suppression improves lifespan and stress resistance. One possibility is that the endogenous decline represents a compensatory response to increasing metabolic and oxidative stress that is insufficient in magnitude or timing to produce the phenotype achieved by sustained RNAi. Alternatively, constitutive neuronal knockdown may alter the developmental or temporal trajectory of dENL/AF9 activity in a manner that differs from its gradual decline during normal aging. A third possibility is that aged neurons become increasingly sensitive to changes in *dENL/AF9* as transcriptional, metabolic, and redox buffering capacities deteriorate. These models remain speculative, but each is consistent with the marked age dependence of the transcriptomic response and emphasizes that both the timing and magnitude of *dENL/AF9* reduction are likely to influence its biological consequences.

The transcriptional response also suggests that *dENL/AF9* suppression engages stress- and translation-associated programs differently across the lifespan. In young flies, the most prominent pathway-level response was negative enrichment of ribosome-associated processes. This observation is notable in relation to our recent neuronal study of the *Drosophila* YEATS2 orthologue, in which *dYEATS2* depletion produced pronounced suppression of ribosomal and translation-associated genes together with alterations in fatty-acid-oxidation and branched-chain-amino-acid-degradation programs [11]. Although ENL/AF9 and YEATS2 participate in distinct chromatin-associated complexes, the recurrence of ribosomal and metabolic signatures raises the possibility that regulation of translational and metabolic adaptation represents a broader feature of neuronal YEATS-domain function. This interpretation is also compatible with our pharmacological ENL/AF9 study [12], in which YEATS-domain inhibition modulated the integrated stress response, a pathway that coordinates translation with nutrient availability, amino-acid balance, oxidative stress, and mitochondrial dysfunction. The present data do not establish direct engagement of the integrated stress response, but they reinforce the concept that YEATS-domain proteins may influence the ability of neurons to adapt transcriptional and translational programs to changes in cellular metabolic state.

The longevity-associated features observed at D40 further argue against a simple canonical insulin/insulin-like signalling mechanism. The negatively enriched KEGG longevity-regulating pathway was driven predominantly by reduced Hsp70-family genes rather than by coordinated changes in classical IIS components, consistent with our previous observation that representative IIS genes were not significantly suppressed at earlier ages [16]. Because Hsp70-family expression commonly increases with age and cellular stress [36, 37], its reduction may reflect an altered basal stress state rather than a pro-longevity effect of reduced chaperone activity itself. Conversely, the proteomic dataset contained Hsp22, Prx2, and *stunted (sun)* within the GO term “determination of adult lifespan.” Increased Hsp22 has been associated with lifespan extension, oxidative-stress resistance, and mitochondrial maintenance [33], while neuronal Prx2/Jafrac1 expression promotes longevity and preserves mitochondrial function [34]; reduced *sun* activity has likewise been linked to extended lifespan and enhanced oxidative-stress resistance [35]. These associations do not establish that any of these proteins mediates the lifespan effect of *dENL/AF9* suppression, but their emergence within the D40 proteomic signature provides an additional connection between the observed metabolic/redox state and established longevity-associated processes. Together with our previous evidence of enhanced catalase activity and reduced lipid peroxidation [16], the findings support a model in which *dENL/AF9* suppression affects metabolic and stress-response plasticity rather than acting primarily through a single canonical longevity pathway.

The matched lipidome–metabolome integration provides an additional level of evidence by asking whether two molecular layers measured in the same biological specimens contribute to a common genotype-associated pattern. The DIABLO model further indicated that the genotype-associated lipid and metabolite changes were organized within a coordinated cross-block signature across the same D40 biological samples. Within this joint structure, the inverse associations between citrate and several triglyceride- and ceramide-associated species suggest coordinated variation between central-carbon and lipid-associated metabolism in *dENL/AF9*-knockdown heads. This relationship is biologically plausible because citrate occupies a key position at the interface between mitochondrial carbon metabolism and cytosolic lipid biosynthesis [38]. However, the present data do not establish whether altered citrate abundance reflects changes in TCA-cycle activity, mitochondrial export, downstream utilization, or carbon partitioning, nor do they demonstrate direct biochemical coupling between citrate and the associated lipid species. The integration should therefore be viewed as evidence of coordinated molecular variation rather than metabolic flux or causal regulation. Given the small matched sample size and the supervised nature of DIABLO, these associations remain hypothesis-generating and are best interpreted together with the single-omics findings.

Several limitations define the scope of these conclusions. Only transcriptomic profiling included both young and aged animals, precluding inference of age dependence for the lipidomic, metabolomic, proteomic, ATP, and NAD/NADH phenotypes. In addition, neuronal *dENL/AF9* knockdown was analyzed using whole heads; consequently, the measured molecular changes may include both neuron-autonomous effects and secondary responses in glial or other head tissues. Metabolite identities remain putative because they were derived from untargeted LC–MS annotation without confirmation against authentic standards, and the pathway-level metabolomic findings should therefore be regarded as candidate metabolic patterns. Although the combined datasets are consistent with altered lipid utilization, central-carbon metabolism, mitochondrial energy-associated processes, and redox balance, they do not establish increased β-oxidation, TCA-cycle activity, oxidative phosphorylation, or respiratory flux. Increased ATP abundance and the elevated NAD/NADH ratio therefore indicate an altered whole-head energetic and redox state rather than direct evidence of enhanced mitochondrial activity. Future targeted metabolite validation, isotope-tracing or respiratory analyses, cell-type-resolved profiling, and direct interrogation of *dENL/AF9*-dependent chromatin regulation will be required to determine how this molecular state is generated and which components contribute causally to the longevity phenotype.

Overall, neuronal *dENL/AF9* suppression elicits a substantially stronger transcriptional response in aged than young fly heads and is associated with a distinct metabolic and energetic molecular state at D40. Cross-omics analysis linked changes in lipid handling and central-carbon and nucleotide metabolism to mitochondrial ATP-related processes. These changes were accompanied by increased ATP abundance and a more oxidized NAD redox state. In addition, matched lipidome–metabolome analysis connected the putatively annotated citrate signal with altered lipid features. These findings extend the physiological role of *dENL/AF9* beyond its previously established association with lifespan and oxidative-stress resilience and provide a systems-level framework for investigating how a YEATS-domain histone-acylation reader intersects with metabolic and stress-response adaptation in the aging nervous system.

## Statements and Declarations

## Acknowledgments

The authors acknowledge the MedCMU research infrastructure center in the Faculty of Medicine, Chiang Mai University. We thank Mr. Papon Hitmool, Mr. Sakorn Phongchankhiao, and Mr. Tawan Munwanna (Department of Pharmacology, Faculty of Medicine, Chiang Mai University) for their technical and administrative assistance.

## Data availability

RNA-seq data from young (D3) flies are publicly available in the NCBI Gene Expression Omnibus (GEO) under accession GSE307078 (BioProject PRJNA1314219). RNA-seq data from aged (D40) flies have been deposited in GEO under accession GSE348592 and will be released publicly upon publication of the peer-reviewed article. Additional information may be obtained from the first or corresponding author upon reasonable request.

## Code availability

Custom Python and R scripts generated for this study are available from the first author or corresponding author upon reasonable request. Data analyses performed using commercially available software and standard analysis tools are described in the Methods.

## Ethics Approval

The Ethics Committee of the Faculty of Medicine, Chiang Mai University, Thailand, approved the experimental protocol in this study (protocol code: 2569/INS-11).

## Materials availability

*Drosophila* stocks used in this study are available from the Bloomington Drosophila Stock Center or from the corresponding author upon reasonable request.

## Authors’ contributions

Conceptualization: L.LP., R.Y.; Data curation: R.Y., P.N., S.D., and L.LP.; Formal Analysis: R.Y., P.N., and S.D.; Funding acquisition; L.LP.; Investigation: R.Y., P.N., S.D., C.P., P.P., S.J., and L.LP.; Methodology: R.Y., P.N.; Project administration: L.LP.; Resources; L.LP, S.J; Software: R.Y., P.N., S.D.; Supervision; L.LP; Validation: R.Y., P.N., L.LP.; Visualization: R.Y., P.N., L.LP.; Writing – original draft: R.Y.; Writing – review & editing: All authors have reviewed and approved the final version of this manuscript.

## Declaration of generative ai and ai-assisted technologies in the writing process

During the preparation of this work, the authors used Claude and ChatGPT (GPT-5.6) to assist with improving the clarity, readability, and organization of the manuscript and with the development and generation of the graphical abstract. Following the use of these tools, the authors critically reviewed, revised, and approved all AI-assisted content and take full responsibility for the final content of the article.

