## Supplementary Figures 1-9 for "Multi-omics profiling identifies a metabolic and energetic signature associated with neuronal *dENL/AF9* suppression in aged flies"

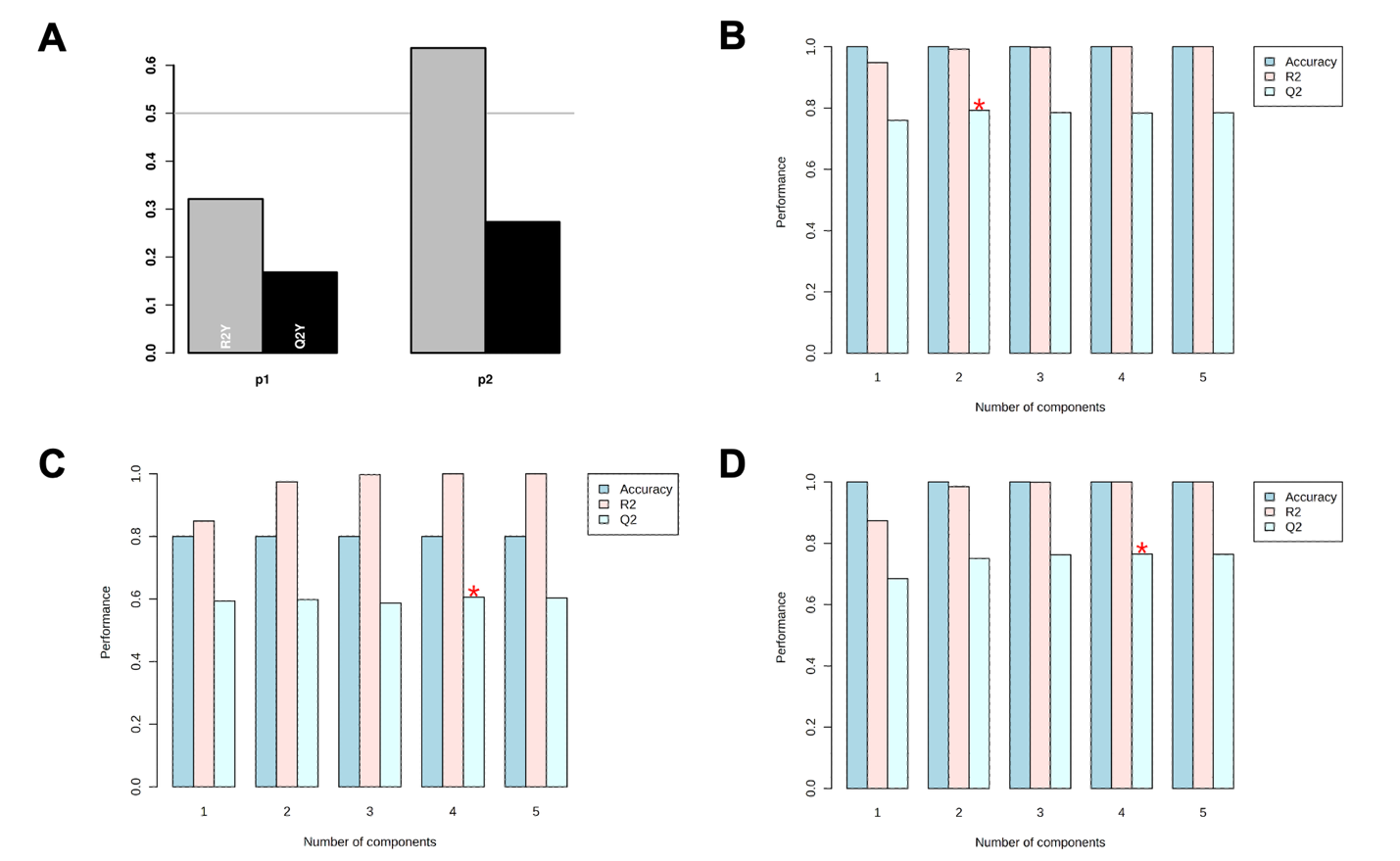

**Supplementary Fig. 1 | Cross-validation of PLS-DA models for transcriptomic, metabolomic, proteomic, and lipidomic datasets.** Cross-validation was used to assess the performance of partial least-squares discriminant analysis (PLS-DA) models distinguishing NTC IR and dENL/AF9 IR fly heads. **(A)** Transcriptomic PLS-DA model performance for the first (p1) and second (p2) latent components, expressed as explained variation (R²Y) and cross-validated predictive ability (Q²Y). For the two-component model, R²Y = 0.636 and Q²Y = 0.273. (B–D) Cross-validation performance of the **(B)** lipidomic**,** **(C)** metabolomic, and **(D)** proteomic PLS-DA models across increasing numbers of latent components. Bars indicate classification accuracy, coefficient of determination (R²), and cross-validated predictive ability (Q²). Red asterisks indicate the number of components selected as optimal by the cross-validation procedure: two components for metabolomics and four components for proteomics and lipidomics.

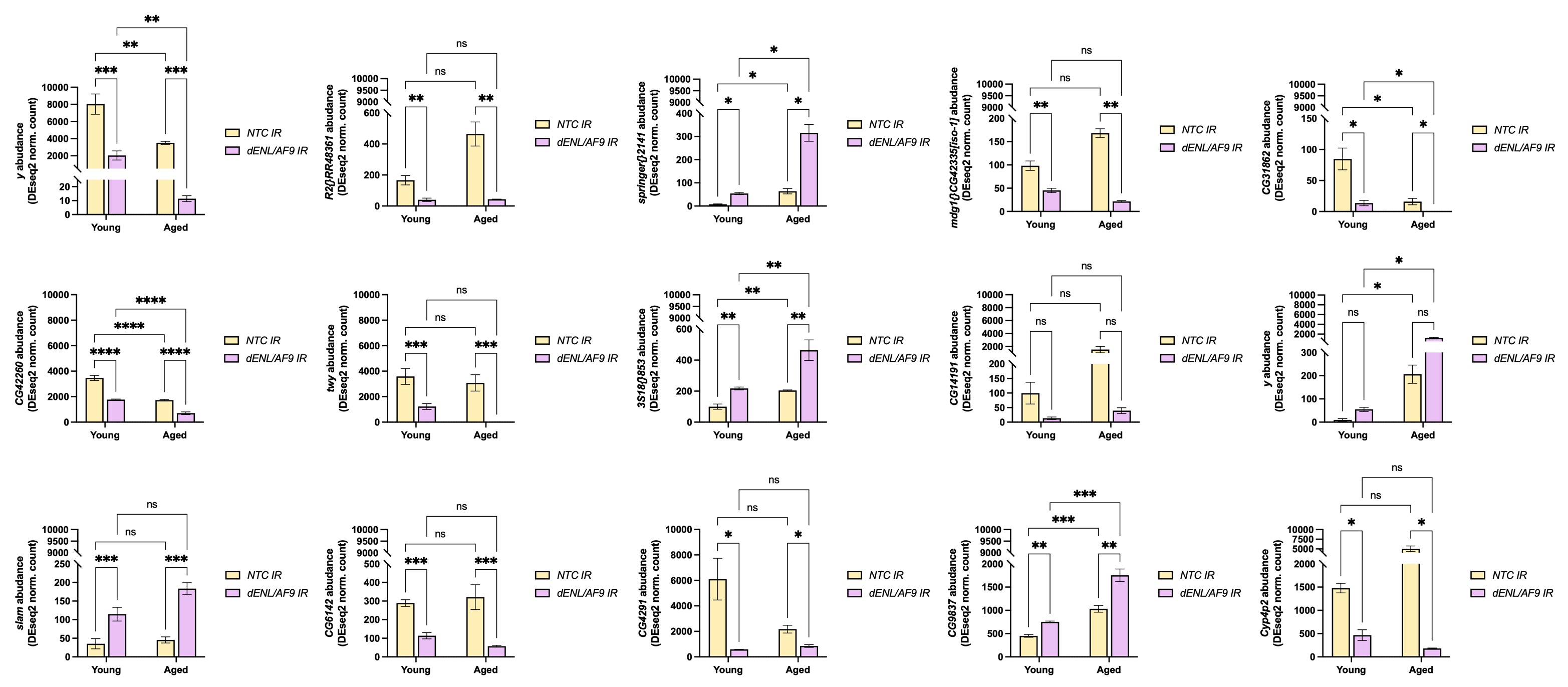

**Supplementary Fig. 2 | Expression patterns of DEGs shared between young and aged *dENL/AF9*-knockdown fly heads.** Normalized abundance of the 15 differentially expressed genes (DEGs) commonly detected in both young and aged transcriptomic datasets. Each plot shows gene expression in NTC IR and dENL/AF9 IR fly heads at the young and aged stages, allowing comparison of age-dependent and genotype-dependent expression patterns. A two-way ANOVA with Tukey's multiple comparisons test was used for the comparisons. Data are shown as mean ± SEM. Statistical significance is indicated as * p<0.05, ** p<0.01, *** p<0.001, **** p<0.0001, and ns = not significant.

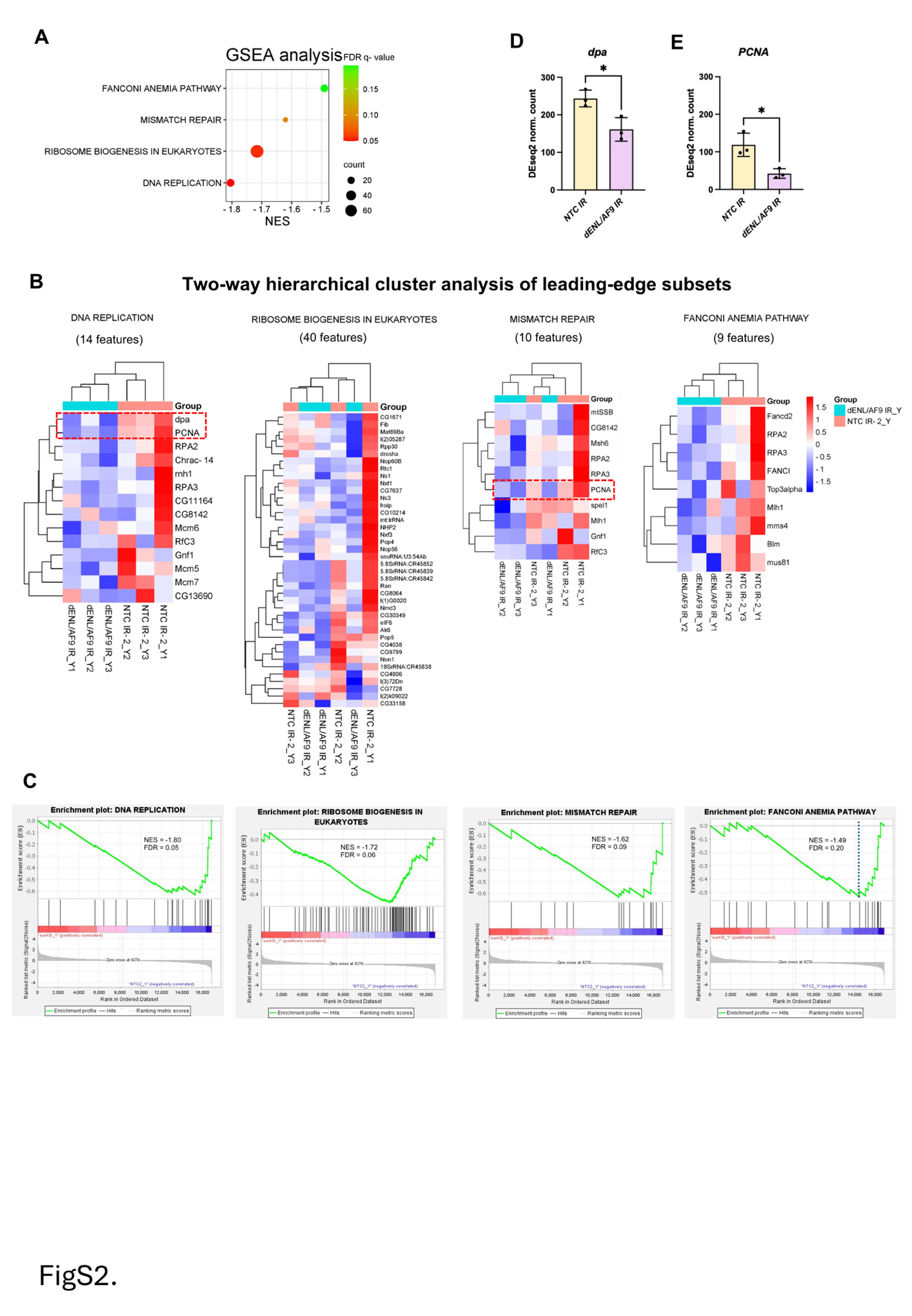

**Supplementary Fig. 3 | Gene set enrichment analysis of young *dENL/AF9*-knockdown fly heads. (A)** Gene set enrichment analysis (GSEA) of transcriptomic data from young *dENL/AF9-*knockdown fly heads relative to age-matched *NTC IR* controls. Dot size indicates gene count, and color indicates FDR q-value. **(B)** Two-way hierarchical clustering heatmaps of the leading-edge subsets from the enriched pathways shown in (A). Genes consistently expressed across all replicates are highlighted in the dotted square. **(C)** Enrichment plots for the four pathways identified by GSEA in young *dENL/AF9*-knockdown fly heads. (**D)** and **(E)** Normalized expression levels of representative genes that are highlighted in (B), in young fly heads. An unpaired t-test was used for the comparisons. Data are shown as mean ± SEM. Statistical significance is indicated as * p<0.05

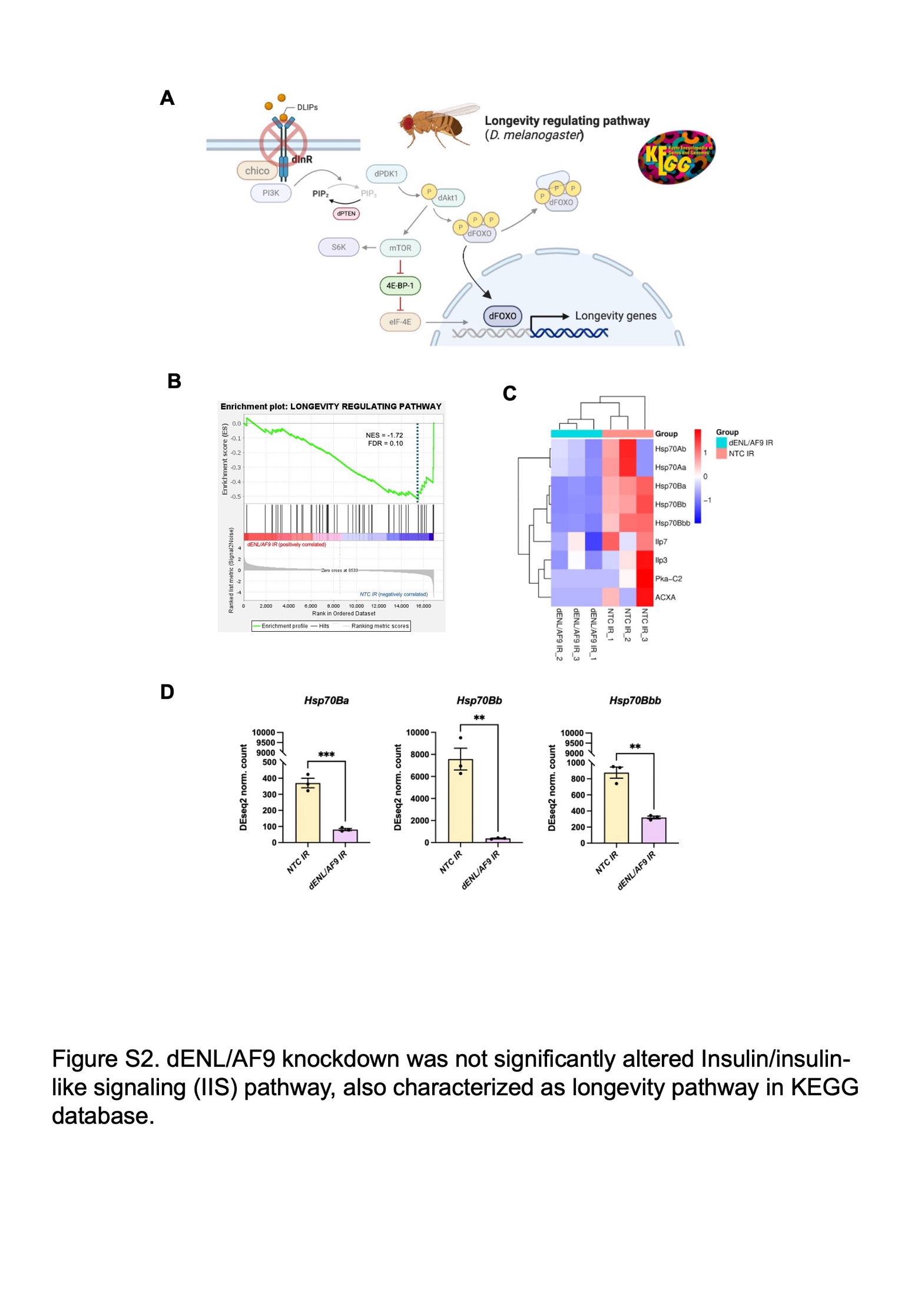

**Supplementary Fig. 4 | Negative enrichment of the longevity-regulating pathway in aged *dENL/AF9*-knockdown heads is driven predominantly by Hsp70-family genes rather than canonical IIS components. (A)** Schematic overview of the insulin/insulin-like signalling (IIS) pathway, annotated as the longevity-regulating pathway (*D. melanogaster*) in the KEGG database. Core components shown include dInR, chico, PI3K, dPDK1, dAkt1, dFOXO, mTOR, S6K, 4E-BP1 and eIF-4E. **(B)** Gene set enrichment analysis (GSEA) plot for the longevity-regulating pathway in aged dENL/AF9-knockdown fly heads relative to age-matched NTC IR controls (NES = −1.72, FDR = 0.10). **(C)** Clustering heatmap of leading-edge subsets from the longevity-regulating pathway identified by GSEA analysis. **(D)** Normalized expression levels of representative leading-edge genes showing consistent genotype-associated expression patterns across biological replicates. An unpaired t-test was used for the comparisons. Data are shown as mean ± SEM. Statistical significance is indicated as * p<0.05,** p<0.01, and *** p<0.001.

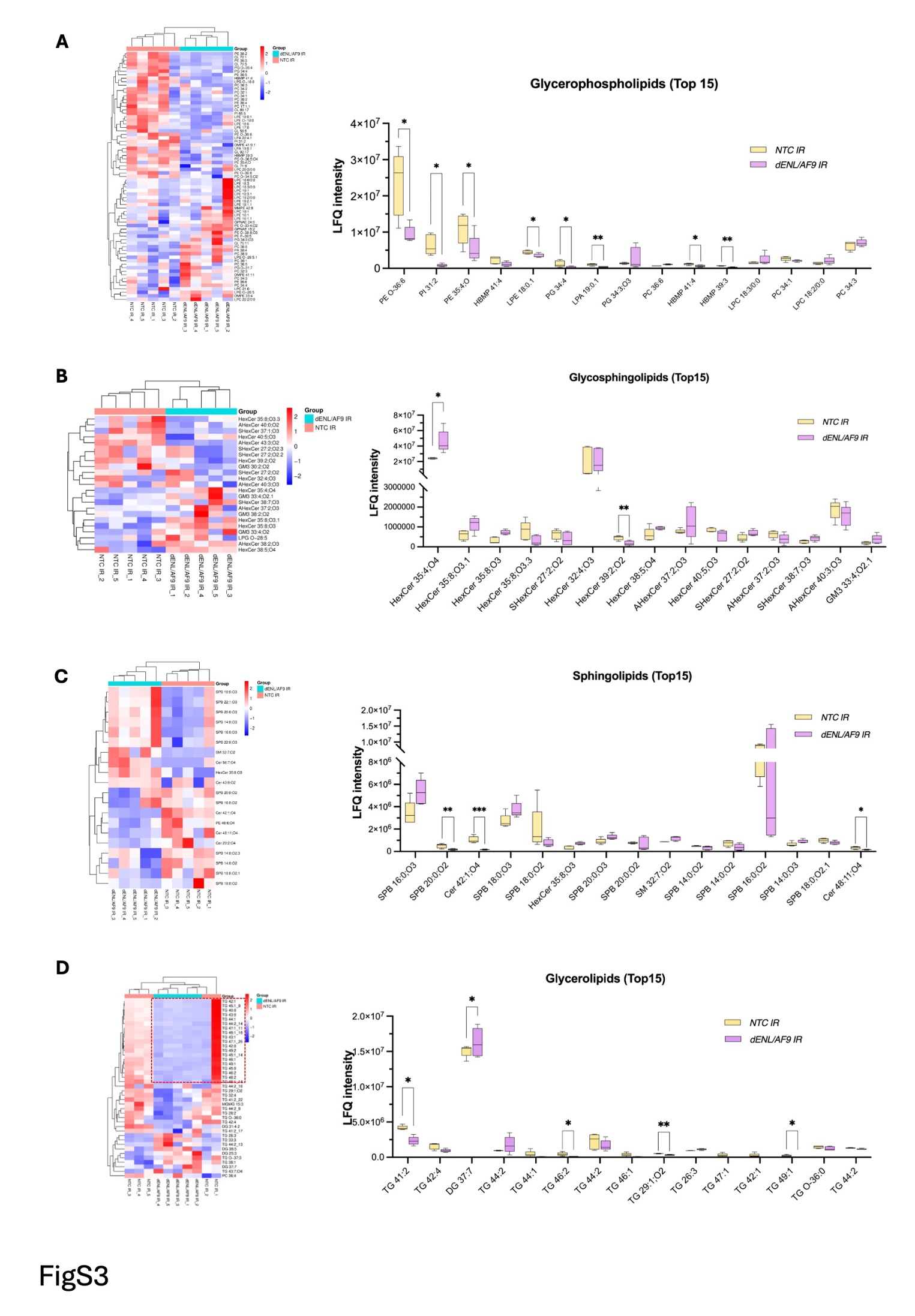

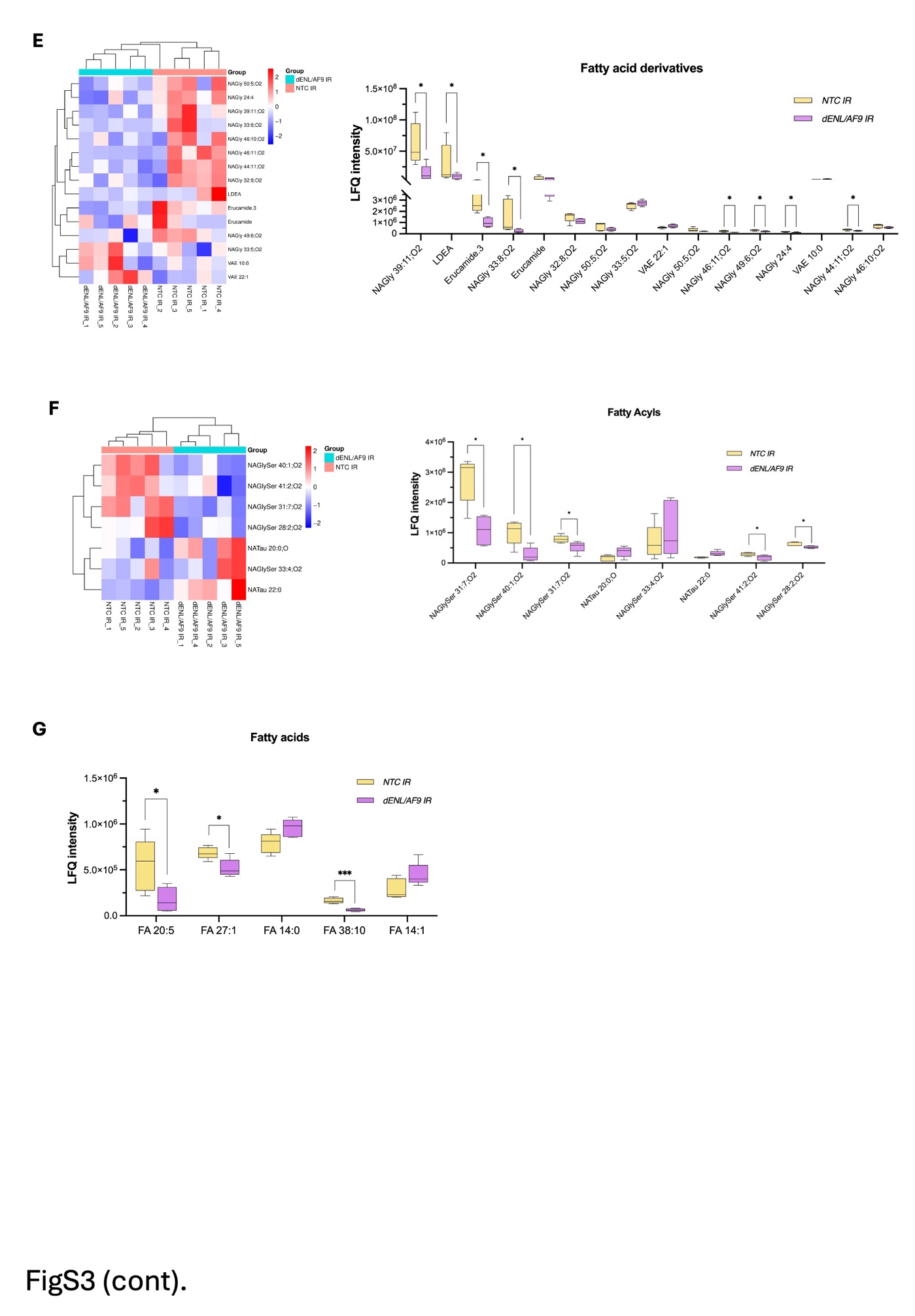

**Supplementary Fig. 5 | VIP lipid categories identified in aged *dENL/AF9*-knockdown fly heads. (A-F)** Hierarchical clustering summarizes category-specific VIP features; the adjacent plots display up to the top 15 features for visualization, and statistically altered features were used to select the species shown in the main figure. Shown are glycerophospholipids (A), glycosphingolipids (B), sphingolipids (C), glycerolipids (D), fatty acid derivatives (E), and fatty acyls (F). For each class, the left panel shows the clustering heatmap of the selected VIP features and the right panel shows the box plots of the relative intensity of the corresponding lipid species. **(G)** Relative abundance of selected fatty acids in aged *NTC IR* and *dENL/AF9 IR* fly heads. Apart from these features, L-arginine was also detected among the lipidomics-derived VIP features. A multiple unpaired t-test with a Bonferroni correction was used for the comparisons. Statistical significance is indicated as * p<0.05, ** p<0.01, and *** p<0.001.

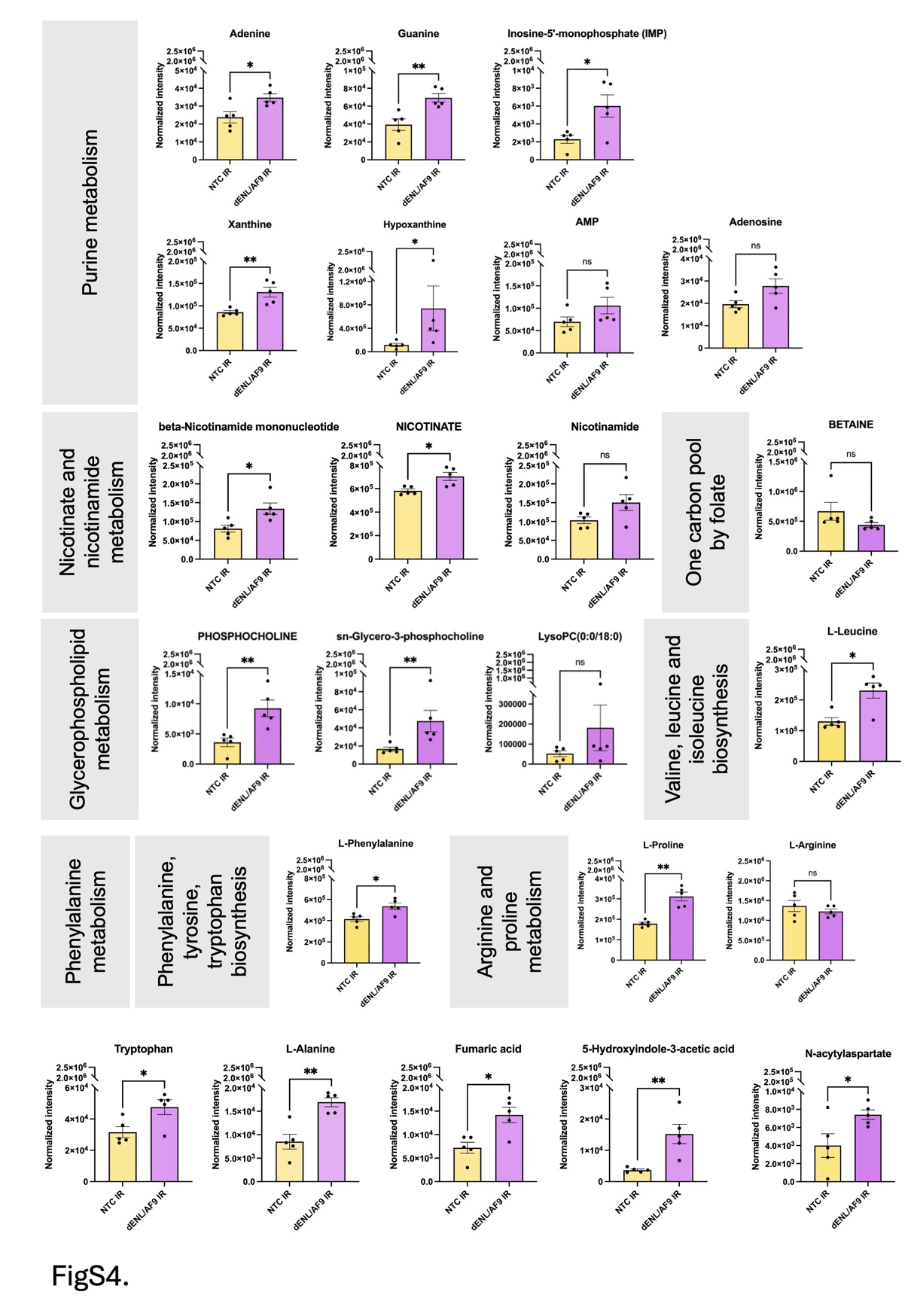

**Supplementary Fig. 6 | Relative abundance of metabolic pathways highlighted by exploratory pathway analysis in aged *dENL/AF9*-knockdown fly heads.** Bar plots show the distribution of normalized feature intensity for VIP metabolites assigned to the pathways identified by metabolomic pathway analysis (Fig. 3E) in aged *NTC IR* and *dENL/AF9* IR fly heads. Additional individual metabolites linked to the integrated metabolic signature are also shown. An unpaired t-test was used for the comparisons. Data are shown as mean ± **SEM.** Statistical significance is indicated as * p<0.05, ** p<0.01, and ns = not significant.

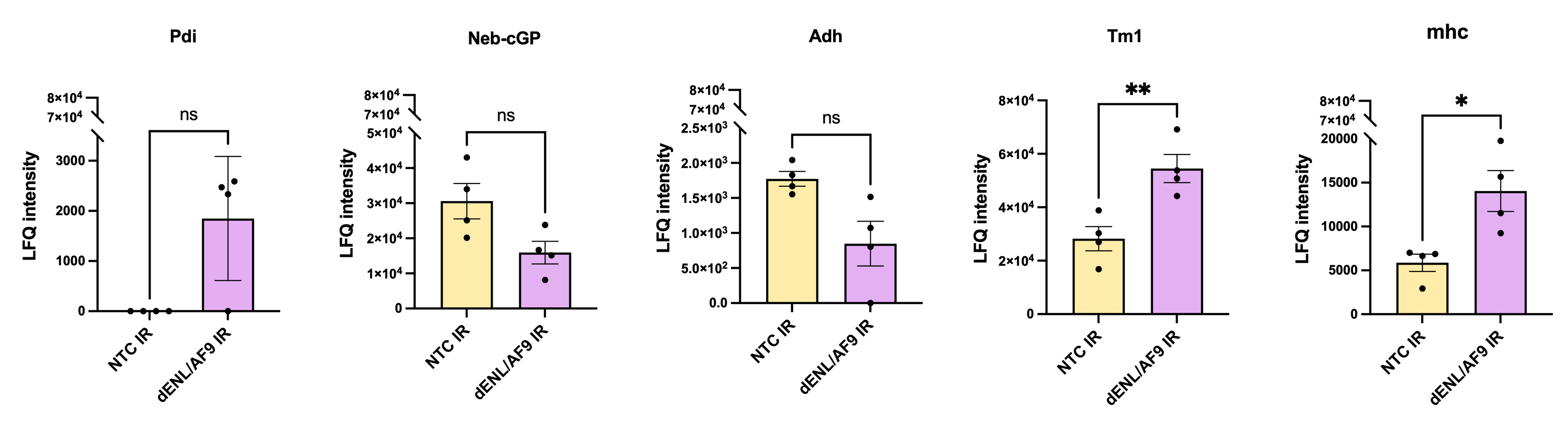

**Supplementary Fig. 7 | Relative abundance of differentially abundant proteins assigned to STRING cluster 1 in aged *dENL/AF9*-knockdown fly heads.** Box plots show the distribution of normalized feature intensity for representative differentially abundant proteins (DAPs) assigned to STRING cluster 1 in aged *NTC IR* and *dENL/AF9 IR* fly heads. An unpaired t-test was used for the comparisons. Data are shown as mean ± **SEM.** Statistical significance is indicated as * p<0.05, ** p<0.01, and ns = not significant.

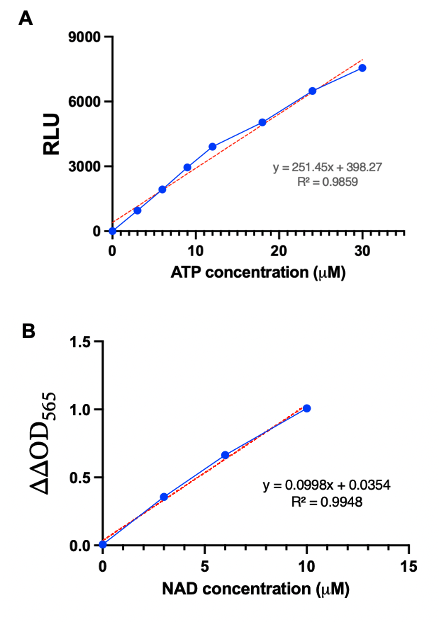

**Supplementary Fig. 8 | Standard curves for ATP and NAD measurement.** Calibration curves used to quantify adenosine triphosphate (ATP) and nicotinamide adenine dinucleotide (NAD) concentrations from assay signals (Fig. 4H and 4I). **(A)** ATP standard curve showing relative luminescence units (RLU) plotted against ATP concentration in micromolar (µM). **(B)** NAD standard curve showing change in absorbance at 565 nm (ΔΔOD565) plotted against NAD concentration in micromolar (µM). Blue dots represent either ATP or NAD standards at each concentration; the blue line shows the fitted calibration curve, and the red dashed line indicates the linear regression. The regression equation and coefficient of determination (R²) are shown on the plot.

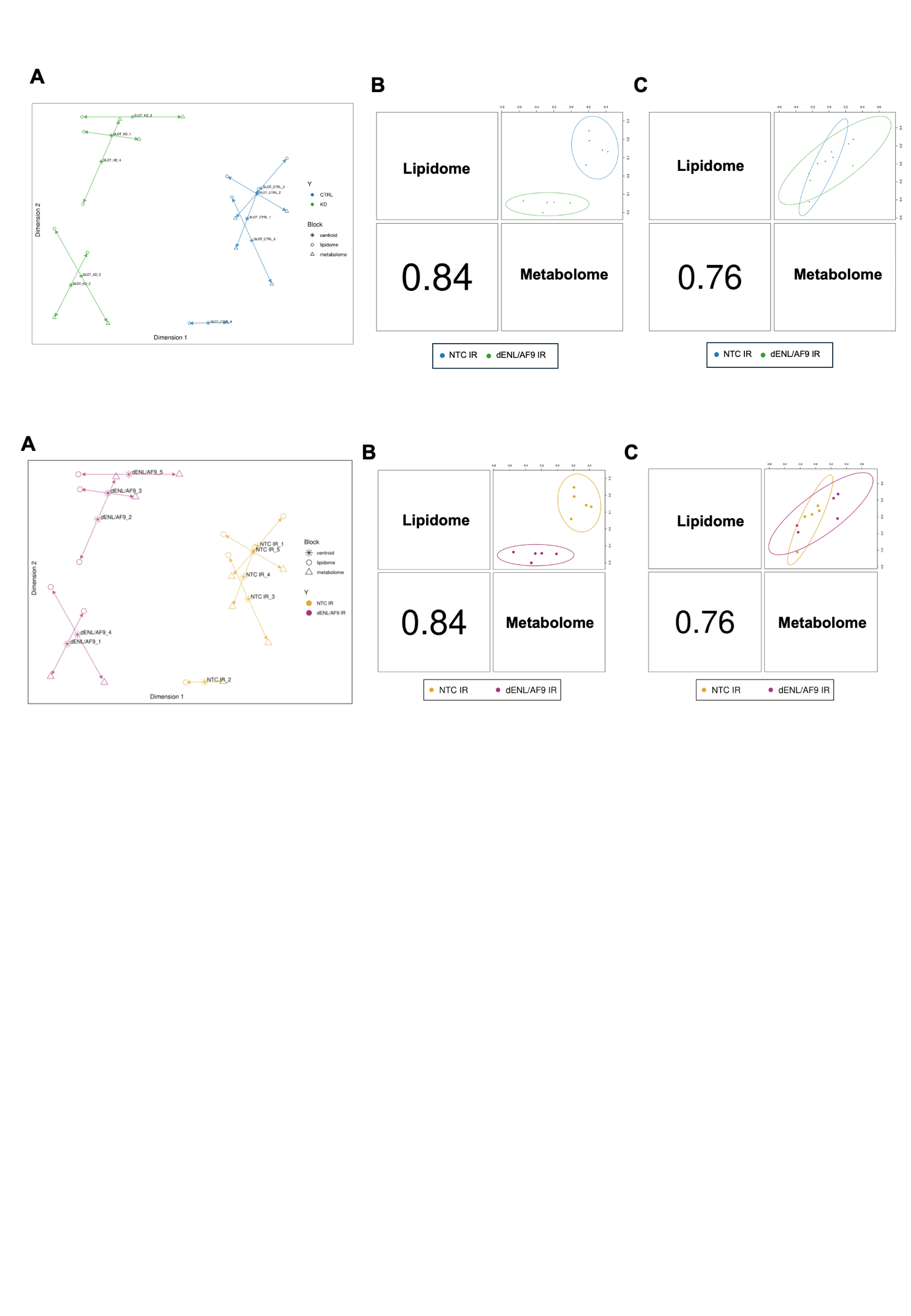

**Supplementary Fig. 9 | DIABLO (Multi-block sPLSDA) integrating multi-omics data.**
(A) DIABLO arrow plot showing block-specific projections in the integrated latent-variable space. Arrows connect the lipidomic and metabolomic projections obtained from the same biological sample. (B) and (C) Pairwise correlations among block-specific latent components for the first (B) and second (C) latent dimensions. Values indicate correlation coefficients between omics blocks within the fitted exploratory model.
